# Location-Specific Facilitation in Primate Auditory Cortex

**DOI:** 10.1101/2022.06.19.496736

**Authors:** Chenggang Chen, Sheng Xu, Yunyan Wang, Xiaoqin Wang

## Abstract

Neural responses to sensory stimuli are markedly influenced by the context in which a stimulus is preceded or embedded. Cortical and subcortical neurons typically exhibit adaptation to repetitive auditory, visual, somatosensory, and olfactory stimulation. Here, we investigated single neuron responses to sequences of sounds either repeatedly delivered from a single spatial location or randomly delivered from multiple spatial locations in the auditory cortex of awake marmosets. Instead of inducing adaptation, repetitive stimulation from a target speaker evoked long-lasting, location-specific facilitation (LSF) in many neurons, irrespective of the visibility of the target speaker. The extent of LSF decreased with decreasing presentation probability of the target speaker. Intracellular recordings showed that repetitive sound stimulation evoked sustained membrane potential depolarization which gave rise to firing rate facilitation. Computational models suggest two distinct neural mechanisms underlying LSF. Our findings revealed a novel form of contextual modulation in the auditory cortex that may play a role in auditory streaming and predictive coding.

## INTRODUCTION

It has been well established that responses of neurons in sensory cortical and subcortical areas are influenced by stimulus context. Contextual effects are mainly suppressive when the same stimulus is repeatedly presented. For example, the presence of a preceding sound suppressed a neuron’s responses to a succeeding sound of the same feature in ascending auditory pathway (Shore, 1995; Nelson et al., 2009; Xiong et al., 2020; Bartlett and Wang, 2005); visual cortical neurons exhibited repetition suppression to the image that is presented twice (Muller et al., 1999; Williams and Olson, 2022); repetitive whisker stimulation reduced the responses of somatosensory neurons (Ganmor et al., 2010; Heiss et al., 2008); and neural activities in the olfactory pathway habituated to the repeated odor presentation (Wilson, 1998; Kato et al., 2012). In addition to temporal context, spatial context can also suppress neural responses in different modalities. For example, along the ascending visual pathway, neural responses to a stimulus at its receptive field are suppressed by the same surrounding stimulus (Huang et al., 2019; Alitto and Usrey, 2008; Keller et al., 2020); somatosensory cortical neural responses to deflection of its principal whisker are suppressed when its neighboring whiskers are deflected (Mirabella et al., 2001; Petersen et al., 2009); and responses of auditory neurons to a probe sound from one spatial location are suppressed by masker sounds from other locations (Fitzpatrick et al., 1999; Reale and Brugge, 2000; Mickey and Middlebrooks, 2005). In a study of awake marmoset auditory cortex, it was found that a masker placed far away from a neuron’s spatial receptive field (SRF) suppressed the response elicited by a probe sound in its SRF, suggesting contextual modulations are widespread and suppressive in the spatial domain (Zhou and Wang, 2012, 2014). The above-cited previous studies suggest that neural adaptation to repetitive stimuli is a general rule in cortical and subcortical neurons across sensory modalities.

An important property of neural adaption in sensory neurons is stimulus-specific adaptation (SSA) to the stimuli that are presented with a high probability. SSA has been studied extensively in different modalities, including auditory (Ulanovsky et al., 2003; Anderson et al., 2009; Malmierca et al., 2009), visual (Kohn and Movshon, 2003; Movshon and Lennie, 1979), somatosensory (Katz et al., 2006; Liu et al., 2017) and olfactory (Wilson, 1998; Verhagen et al., 2007) systems. SSA has attracted much interest in the past several decades because it was thought to be a potential neuronal correlate of mismatch negativity (MMN) (Ulanovsky et al., 2003; Fishman and Steinschneider, 2012), which has been extensively studied in humans (Näätänen et al., 2007). SSA is also believed to reflect deviance detection (Polterovich et al., 2018; Pérez-González et al., 2021), predictive coding (Parras et al., 2017; Carbajal and Malmierca, 2018) and efficient coding (Wark et al., 2007; Kohn, 2007). Visual and somatosensory neurons exhibit SSA in both spatial (Kohn and Movshon, 2003; Liu et al., 2017) or non-spatial (Movshon and Lennie, 1979; Maravall et al., 2007) domains. In contrast, in the auditory and olfactory systems, SSA has been studied mainly by presenting stimuli with two different frequencies (Natan et al., 2015; Kato et al., 2015; Harpaz et al., 2021) or odorants (Kum et al., 2019).

However, the adaptation alone does not account for the ability of sensory systems to remain vigilant to the environment in spite of repetitive sensory stimulation. In this study, we explored spatial contextual modulations by stimulating neurons in the awake marmoset auditory cortex with sequences of sounds either randomly from various spatial locations (equal-probability mode) or repeatedly from a single location (continuous mode). To our surprise, instead of only inducing adaptation as expected from well-documented SSA literature, repetitive stimulation in the high probability mode from spatial locations away from the center of a neuron’s SRF evoked lasting facilitation and adaptation observed by extracellular recordings from single neurons in the auditory cortex. Nearly half of the sampled neuronal population exhibited this spatial facilitation, irrespective of the stimuli type and visibility of the test speaker. The extent of the facilitation decreased with decreasing presentation probability of the test speaker. Intracellular recordings showed that repetitive sound stimulation evoked sustained membrane potential depolarization that was followed by firing rate facilitation. We used computational models to explore neural mechanisms underlying neural facilitation. Taken together, our findings revealed location-specific facilitation (LSF) to repetitively presented sound stimuli in the auditory cortex that has not been observed. This form of spatial contextual modulation may play a role in such functions as detecting the regularity, segregating the sound stream, and solving the cocktail party problem.

## RESULTS

### Repetitive sound stimulation evoked neural facilitation

We evaluated how extracellularly recorded individual neurons in the auditory cortex responded to broadband sounds played from different speaker locations. Fifteen equally spaced speakers were placed on a semi-spherical surface centered around the animal’s head and above the horizontal plane (Figures 1A and 1B). In each test session, we first probed a neuron’s spatial selectivity by delivering a frozen wideband noise stimulus from each speaker location in a randomly shuffled order. We will refer to this stimulation mode as the *equal-probability presentation mode* for which all locations have the same occurrence probability of 1/15. Spatial receptive field (SRF) was constructed for each neuron using the averaged firing rate of the responses to wideband noises in the equal-probability presentation mode. Figures 1C and 1D show the responses of an example neuron obtained in the equal-probability presentation mode. This neuron had an SRF centered around speaker #7 (Figure 1C) and responded to this location with sustained firing throughout the stimulus duration, and to other locations with onset or transient firing (Figure 1D). Figure 2A shows firing rate versus speak number for the equal-probability presentation mode. Speaker #7 evoked the highest firing rate (38.5 spikes/sec) in this neuron, followed by speaker #8 (10.5 spikes/sec). Speakers #14, #13, #9 and #11 evoked near zero firing rate, and a negative firing rate was observed for speakers #2, #4, and #5. Because the firing rate for each speaker location is calculated by subtracting the spontaneous firing rate from the total firing rate, a negative firing rate indicates inhibition.

**Figure 1.**
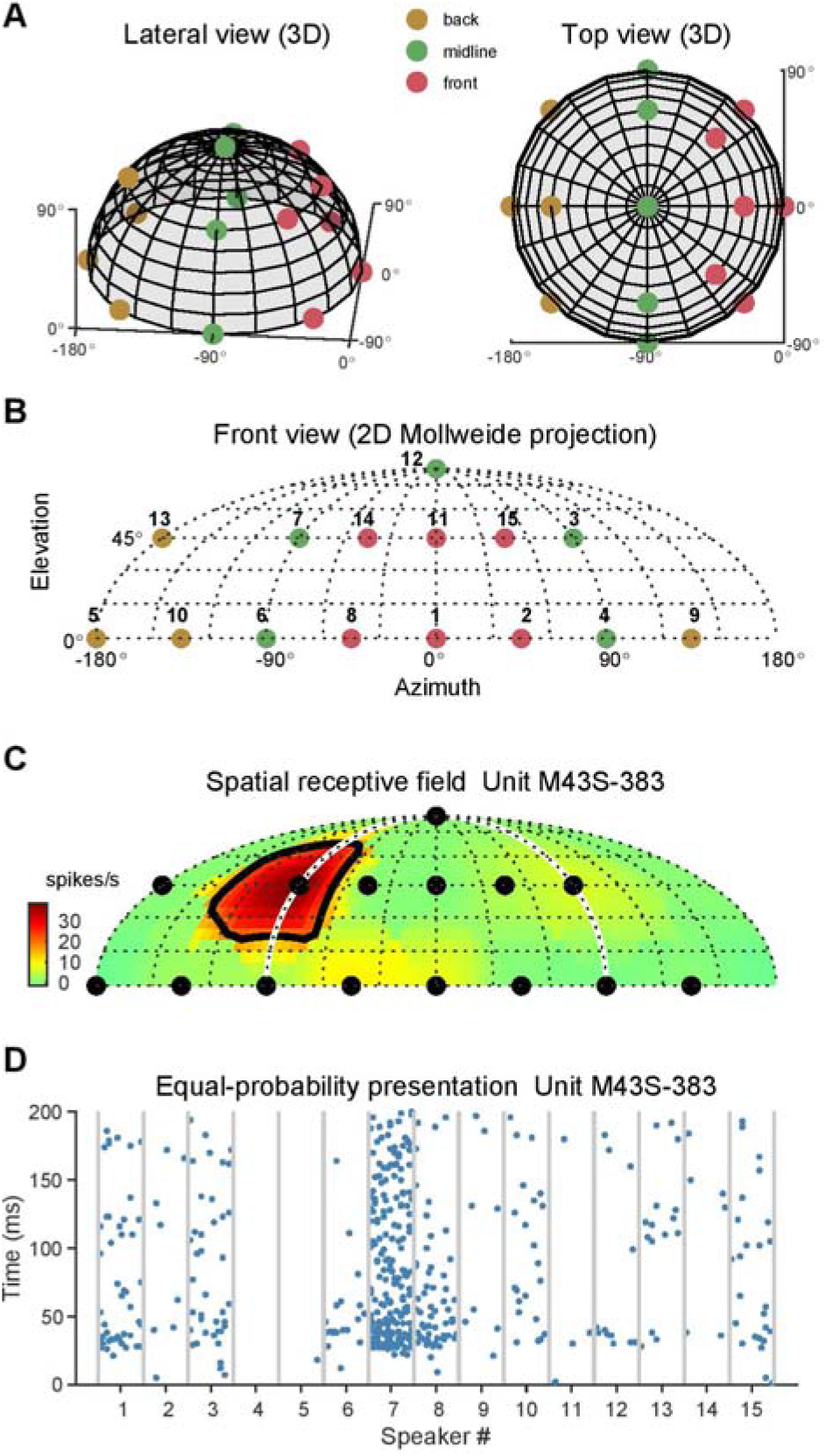
Speaker layout and equal-probability sound stimulation. (A) Fifteen equally spaced speakers were placed on a semi-spherical surface centered around the animal’s head and above the horizontal plane. View from the front, lateral 35° and elevated 30° (left), and view from directly top (right). Red dots indicate six front speakers, green dots indicate five midline speakers, and brown dots indicate four back speakers. (B) Three-dimensional front-back space was projected to a two-dimensional plane around midline for displaying purposes. (C) Spatial receptive field of example Unit M43S-383. White semicircle is the boundary of the front-back space. Black line is the threshold which is defined as the half-maximum firing rate. Black dots indicate fifteen speaker locations. (D) Spike raster plot of same example neuron at fifteen speaker locations under equal-probability presentation mode. Stimuli at each speaker location were randomly presented ten times.

**Figure 2.**
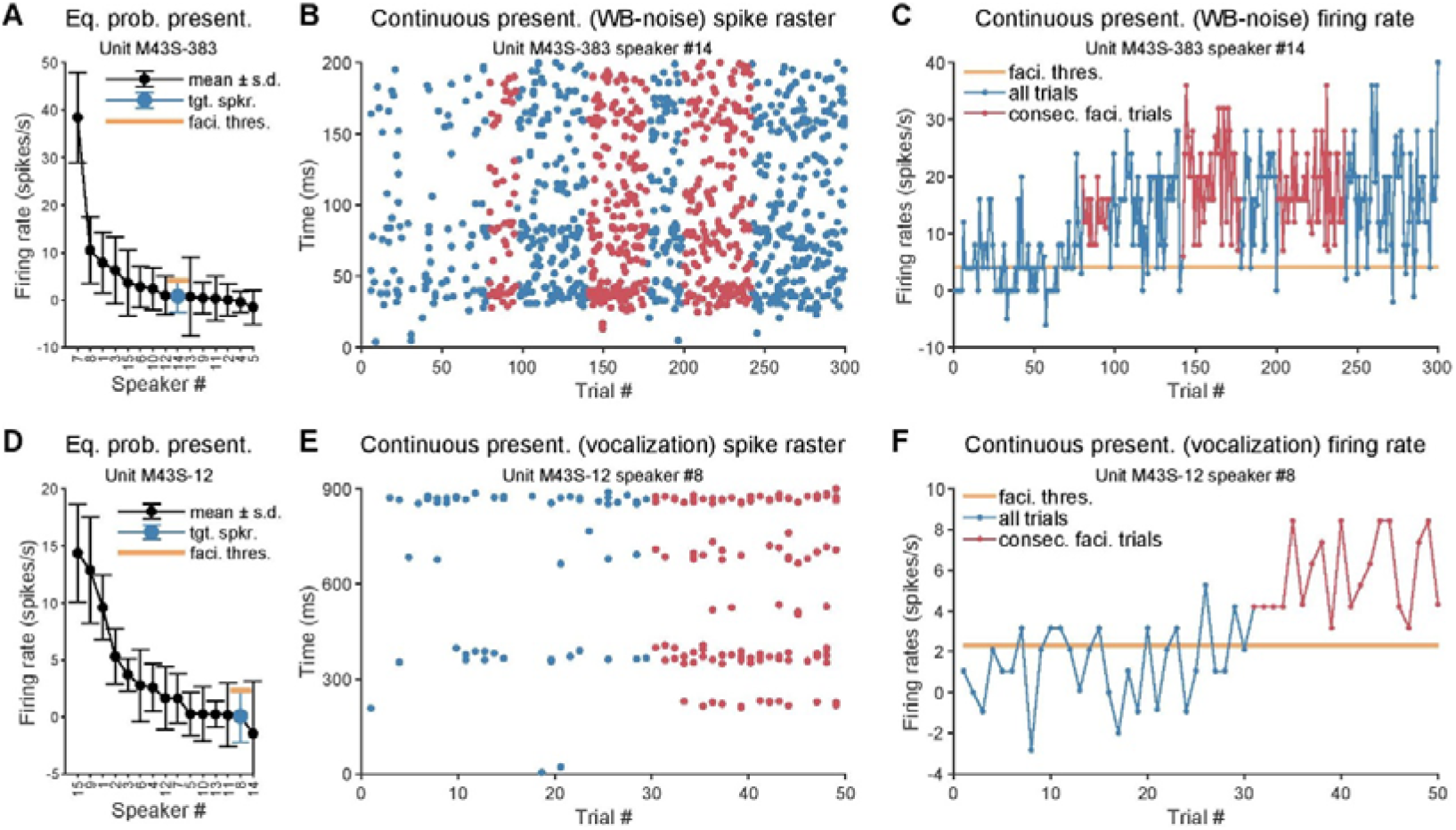
Repetitive sound stimulation evoked neural facilitation. (A) Firing rate versus speak number of unit M43S-383 recorded under the equal-probability presentation mode (same data as Figure 1D). Blue dot and orange bar indicate the target speaker and facilitation threshold (mean + standard deviation), respectively. (B), Spike raster of same example neuron tested at speaker #14 under continuous presentation mode. Red dots indicate spikes belong to the long facilitation phase (i.e., at least five consecutive trials with firing rates exceeding the facilitation threshold). (C), Trial-by-trial firing rate of same example neuron. Red dots and line indicate trails belong to the long facilitation phase. Thick orange line indicates the facilitation threshold. (D-F), Similar to (A-C), but for Unit M43S-12 that was tested using marmoset vocalization stimuli.

After the characterization of a neuron’s SRF, we further tested each neuron with the *continuous presentation mode* in which stimuli were delivered from a speaker location repeatedly, with each trial separated by an inter-stimulus interval of fixed or variable length (range: 500 to 5200 ms, see below). Figures 2B and 2C show the responses of the same neuron depicted in Figures 1C and 1D and Figure 2A to 300 presentations of a 200 ms frozen wideband noise stimulus delivered from speaker #14 in the continuous presentation mode. Speaker #14 evoked 0.8 spikes/sec firing rate in the equal presentation mode (Figure 2A, blue dot) which was considered as the baseline firing rate of this speaker location. In contrast to the expectation from previous auditory cortex literature on adaptation, the response of this neuron to the repeated presentations of a wideband noise stimulus from the same speaker #14 showed epochs consisting of consecutive trials with substantially higher firing rates than the baseline firing rate (median: 16 spikes/sec, Figure 2C, red dots). Elevated firing rates could be observed in trials long after the first trial (e.g., between 150th and 250th trials). Also note that during the trials with elevated firing rates (Figure 2B, red dots), the firing patterns were sustained throughout the stimulus duration. To further examine the facilitated response in the continuous presentation mode, we measured the “facilitation phase” to characterize trials with firing rates exceeding the facilitation threshold which was defined as one standard deviation above the baseline firing rate (Figures 2A, 2C, orange bar). This neuron exhibited several facilitation phases lasting up to 42 trials (Figure 2C, red dots). Figures 2D-2F show responses of another example neuron. In the equal-probability presentation mode, this neuron had weak responses to marmoset vocalization stimuli at speaker #8 (Figure 2D, blue dot, 0.05 spikes/sec). When tested in the continuous presentation mode with speaker #8 (Figures 2E, 2F), this neuron had weak responses initially, but the responses gradually built up and eventually led to a facilitation phase (median: 4.8 spikes/sec) lasting 20 trials (Figure 2F, red dots).

### Neural facilitation occurred in a variety of stimulus conditions

We tested a total of 104 auditory cortex neurons in four hemispheres of three marmosets using wideband stimuli, including frozen wideband noises, amplitude-modulated wideband noises, and marmoset vocalizations. 725 sessions were tested by both equal-probability and continuous presentation modes. Population statistics of the facilitation phase are shown in Figure S1A which shows that a facilitation phase could persist as long as 45 trials. For further analyses, we focused on the sessions that exhibited facilitation phases lasting at least 5 consecutive trials (129 sessions from 51 neurons). In the majority of sessions, 200 trials were tested (Figure S1B). In most sessions, it took fewer than 100 trials to achieve the first facilitation phase lasting at least 5 consecutive trials (median: 44 trials) (Figure S1C).

We calculated the proportion of facilitation trials in the continuous presentation mode for the 129 sessions from 51 neurons that exhibited facilitation phases lasting at least 5 consecutive trials which ranged between 15%-87.5% (median: 32.8%) (Figure 3A, orange line). As a control, we also calculated the proportion of facilitation trials in the equal-probability presentation mode for the same group of 51 neurons (Figure 3A, blue line) which was significantly smaller than that of the continuous presentation mode (14.3% vs. 32.8%, p < 0.0001, rank-sum test). The group of 51 neurons scattered across all recorded cortical areas and did not show any clustering patterns (Figures S2A-S2D). Most of the neurons (43/51) were recorded at superficial cortical depths (< 1 mm, Figures S2E, S2F). We attempted to distinguish putative excitatory and inhibitory neurons by their spike waveform (broad or narrow; Figure S3A) (Mitchell et al., 2007). Spike waveform of 32 neurons had been recorded and 29 neurons had a signal-to-noise ratio higher than 20 dB (Figure S3B). 23 neurons were classified as putative excitatory neurons (yielded 58 sessions) and 6 neurons as putative inhibitory neurons (yielded 12 sessions) (Figure S3C). The proportions of facilitation trials were similar between putative inhibitory and excitatory neurons (Figure S3D; 34% vs. 32%, p = 0.2271, rank-sum test). This analysis suggests that the facilitation can be induced in both putative excitatory and inhibitory neurons.

**Figure 3.**
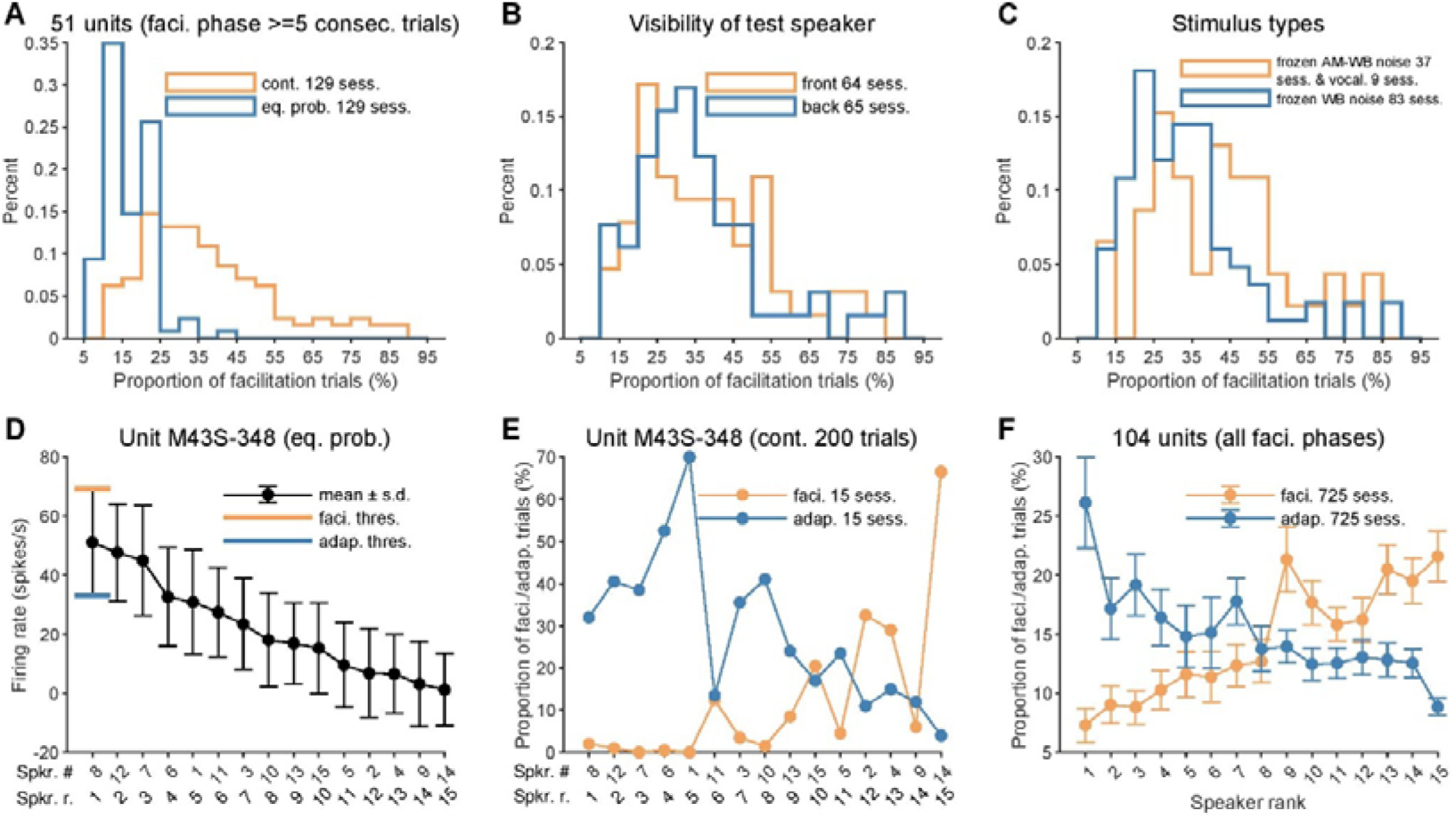
Neural facilitation occurred in a variety of stimulus conditions and depends on sound locations. (A) Histogram of the proportion of facilitation trials under the continuous (orange line) and equal-probability (blue line) presentation modes. Only 129 sessions from fifty-one neurons that exhibited facilitation phases lasting at least five consecutive trials were shown. (B) Histogram of the proportion of facilitation trials under the continuous presentation mode for target speakers located in the front (orange line) and back (blue line). Front speakers: #1, #2, #8, #11, #14, and #15. Back speakers: #3, #4, #5, #6, #7, #9, #10, #12, and #13. (C) Histogram of the proportion of facilitation trials under the continuous presentation mode using frozen amplitude-modulated wide-band noise or animal vocalization (orange line) and frozen unmodulated wide-band noise (blue line) as stimuli. (D) Each speaker was assigned a rank number based on its baseline firing rate obtained under the equal-probability presentation mode. Speaker ranked 1st had the highest firing rate. In this example unit M43S-348, speaker #8 ranked 1st and speaker #14 ranked 15th. Orange and blue bars indicate facilitation and adaptation threshold, respectively. Dots and error bars indicate mean ± standard deviation. (E) Proportion of facilitation (orange dots and lines) and adaptation (blue dots and lines) trials under continuous presentation mode at different speaker ranks for the same example unit. Stimuli at each speaker location were tested 200 times. (F) Proportion of facilitation (orange dots and lines) and adaptation (blue dots and lines) trials of the population data. All 725 sessions from 104 neurons were shown, regardless of the length of the facilitation phase.

We conducted control tests to see if the observed facilitation depended on visual inputs. We found that the facilitation was not limited to speakers located within a marmoset’s visual field (< 90 degrees) (Chaplin et al., 2012) and could be induced from speaker locations both in front and behind an animal (Figures S4A-S4C). Across all 129 test sessions, the proportions of facilitation trials were similar between front (64 sessions) and back (65 sessions) speaker locations (Figure 3B; 38% vs. 35%, p = 0.4653, rank-sum test). The facilitation could still be observed when an animal was tested in the darkness (7 sessions) (Figures S4D-S4F). Interestingly, the proportion of facilitation trials was significantly higher in the darkness than in the light-on condition (122 sessions) (Figure S4G; 57% vs. 32%, p < 0.0001, rank-sum test). These results suggest that visual inputs are not required to induce the facilitation. We also tested the effects of different stimulus types. Neural facilitation was also observed using unfrozen wideband noises (Figure S4H). Comparing to frozen wideband noises (83 sessions), slightly larger proportion of facilitation trials was observed using complex stimuli including amplitude-modulated frozen wideband noises (37 sessions) or marmoset vocalizations (9 sessions) as stimuli (Figure 3C; 30% vs. 41%, p = 0.0054, rank-sum test).

Neural facilitation was observed at both short inter-stimulus intervals (ISI) (700 ms, Figure 2B) and long ISI (1900 ms, Figure 2E). In addition to ISI with a fixed length, we also tested random ISIs in a subset of sessions. An example is shown in Figure S5A. This neuron not only showed facilitation at three constant ISIs but also at random ISIs. Across all 129 test sessions, three groups of ISIs were tested: short (500 ms and 700 ms, 78 sessions), long (>1000 ms, 36 sessions) and random (700 ms to 2200 ms, 15 sessions). The proportion of facilitation trials was similar between the three ISI groups (Figure S5B; 35%, 41%, and 34%, p = 0.1244, one-way ANOVA).

### Neural facilitation did not alter a neuron’s SRF

Previous studies in both auditory and visual cortices found that after presenting a stimulus repeatedly, neurons typically exhibited a decrease of response to the stimulus but an increase of response to other stimuli that are sufficiently different from the repeated stimulus (Condon and Weinberger, 1991; Dragoi et al., 2000), which suggests changes in a neuron’s receptive field. To investigate whether neural facilitation altered a neuron’s SRF, we compared the SRF measured before and after testing a neuron in the continuous presentation mode. An example neuron is shown in Figure S6A (same neuron in Figure 1C). After presenting 300 trials of wideband noise at the same location continuously (spike raster shown in Figure 2B), the SRF (Figure S6A, bottom) appeared similar to the SRF measured before (Figure S6A, top). More examples of pre and post SRFs are shown in Figure S6B. To quantitatively characterize SRF changes, we calculated three metrics in 61 neurons in which pre and post SRFs were measured: tuning selectivity, direction selectivity, and correlation coefficient. The average tuning selectivity (Figure S6C, left, 0.153 vs. 0.146) and direction selectivity (Figure S6C, right, 0.477 vs. 0.473) remained similar after tested by the continuous presentation mode. To further quantify the tuning similarity, we computed the pairwise correlation coefficient between each pair of responses to fifteen speaker locations (Figure S6D, left). 75% (197/262) of pair of sessions had a correlation coefficient greater than 0.7 (Figure S6D, right). We also examined if firing rate changed after a neuron was tested by the continuous presentation mode for the highest (1st) and lowest (15th) ranked target speakers (Figure S6E). Both speaker ranks showed similar firing rates before and after being tested by the continuous presentation mode (ranked 1st: 13.8 vs. 13.0 spikes/sec; ranked 15th: 2.9 vs. 2.6 spikes/sec), consistent with the observations on SRF stability (Figures S6A-S6D). These analyses show that the repetitive presentation of stimuli from the same speaker location does not significantly alter the SRF of the neuron being tested.

In the 129 sessions that exhibited facilitation phases lasting for at least 5 consecutive trials when tested by the continuous presentation mode, the average firing rate during the facilitation phases was nearly three times greater than that evoked by the same speaker location under the equal-probability presentation mode (Figure S6F, orange box: 17.5 vs. 6.6 spikes/sec, p < 0.0001, one-way ANOVA). The spontaneous firing rates during equal-probability presentation mode, non-facilitation and facilitation phases of the continuous presentation mode were similar (Figure S6F, blue box: 5.1, 5.2, and 6.2 spikes/sec, p = 0.3924, one-way ANOVA). Thus, the facilitation phases during the continuous presentation mode were not accompanied by significant changes in spontaneous activities.

### Neural facilitation depends on sound locations

Neural facilitation was observed at all tested speaker locations. To investigate the effect of speaker location on the facilitation, each speaker was assigned a rank number based on its baseline firing rate obtained under the equal-probability presentation mode. Speaker ranked 1st had the highest firing rate among all tested speakers and was at or near the center of a neuron’s SRF. The lowest ranked speaker usually fell far outside of the SRF and evoked a response not significantly different from the spontaneous activity. The speaker rank of an example neuron is shown in Figure 3D. We found that speakers with lower ranks usually elicited more facilitation phases than speakers with higher ranks when tested under the continuous presentation mode as shown by the example neuron in Figure 3E, orange dots. Note that in this neuron no facilitation was induced by the frontal speaker (speaker #1, rank 5th) and only one facilitation trial (out of 200 trials) was induced by the speaker at the contralateral 90° location (speaker #6, rank 4th). These two locations were commonly used in previous SSA studies (see Discussion). When the test speaker was ranked 15th, 18 of 77 (23.4%) tested sessions had facilitation phases lasting for at least 5 consecutive trials (Figure S7A) and the average proportion of facilitation trials was 21.6% (Figure 3F, orange dots). In comparison, when the test speaker was ranked 1st, these statistics dropped to 10.5% (4/38 tested sessions, Figure S7A) and 7.3% (Figure 3F, orange dots), respectively.

In addition to the facilitation, suppressed responses were also observed in the continuous presentation mode. We measured the “adaptation phase” to characterize trials with firing rates lower than the adaptation threshold which was defined as one standard deviation below the baseline firing rate (Figure 3D, blue bar). At higher ranked speakers, more adaptation phases were observed than at lower ranked speakers as shown by an example neuron in Figure 3E, blue dots. Note that 70% of trials (140/200) exhibited adaptation at the frontal speaker (speaker #1, rank 5th) and 52.5% of trials (105/200) showed adaptation at the contralateral 90° location (speaker #6, rank 4th). The average proportion of adaptation trails across all tested sessions was 26.1% when the test speaker was ranked 1st, whereas that number dropped to 8.9% when the test speaker was ranked 15th (Figure 3F, blue dots). This trend was the opposite of facilitation (Figure 3F, orange dots).

### Neural facilitation primarily depends on sound location but not firing rate

We showed above that neural facilitation depends on speaker rank, which was determined by firing rate in the equal-probability presentation mode at a neuron’s preferred sound level. Sound level was a crucial parameter in influencing the firing rate of auditory cortex neurons (Sadagopan and Wang, 2008; Wang, 2018). This gives us the opportunity to determine whether the neural facilitation primarily depends on sound location or any type of stimuli that could influence the firing rate, including speaker location and sound level. Therefore, we measured a neuron’s responses to wideband noise delivered from fifteen speaker locations at four sound levels under the equal-probability presentation mode in a subset of neurons. An example of this analysis is shown in Figure 4A. We assigned this neuron a speaker rank and sound level rank. In contrast to previous figures where the speaker rank was determined by firing rate at one sound level (mostly the best sound level), the speaker rank in this analysis was determined by the averaged firing rate (Figure 4A, black dots) at four sound levels (Figure 4A, four different color dots). The sound level rank at each speaker location was also determined by the firing rate in which the 1st ranked sound level had the highest firing rate among the four tested sound levels. If the neural facilitation was location-dependent, then the facilitation shall increase from higher ranked speakers to lower ranked speakers, but not from higher ranked sound levels to lower ranked sound levels. If the neural facilitation was firing rate-dependent, then the facilitation shall increase from higher ranked speakers to lower ranked speakers as well as from higher ranked sound levels to lower ranked sound levels.

**Figure 4.**
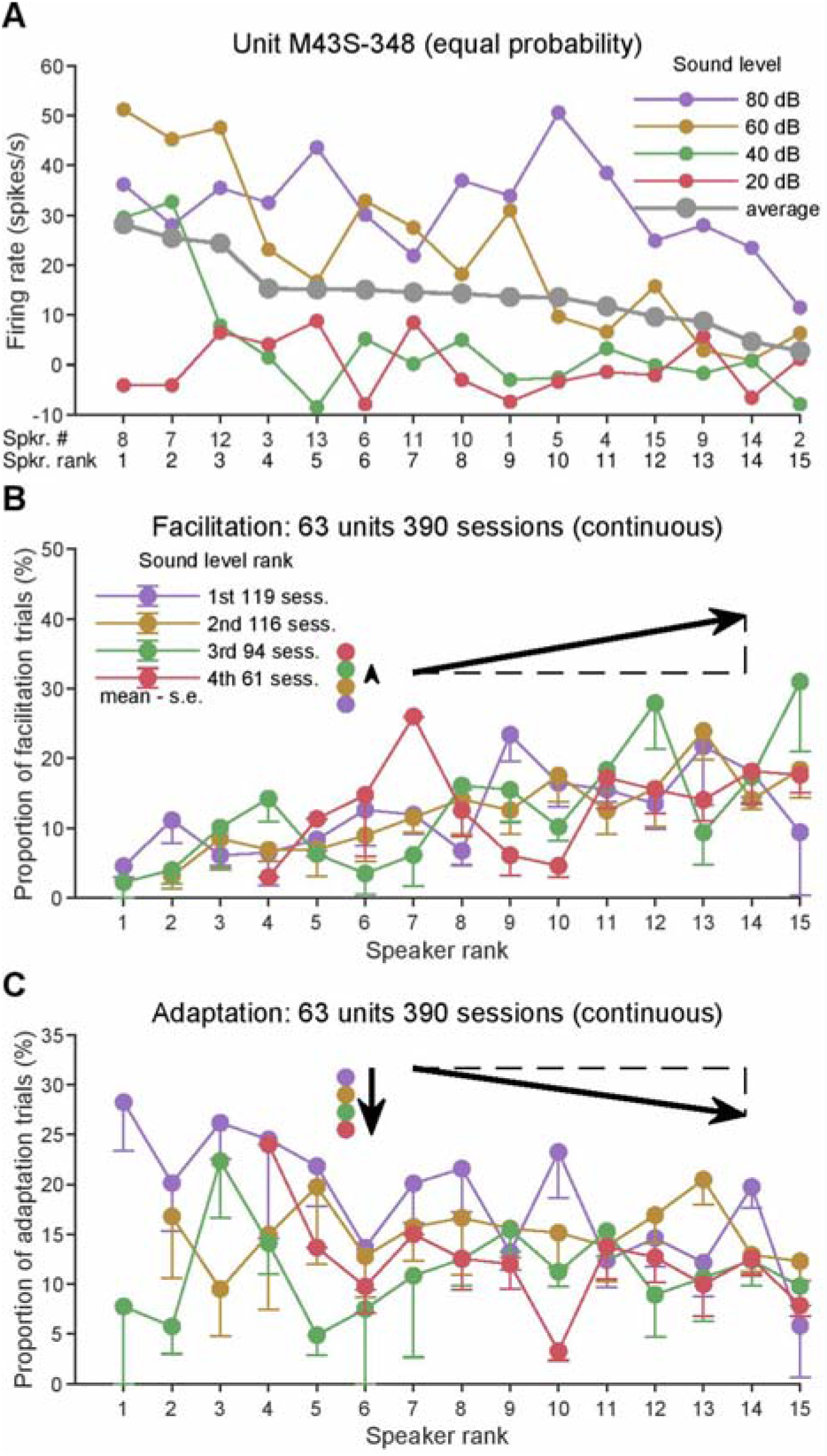
Neural facilitation primarily depends on sound location but not firing rate. (A) Example unit M43S-348’s averaged responses to wide-band noise played at four sound levels across fifteen speaker locations (violet/brown/green/red dots and lines). Speaker rank was determined by the averaged firing rate at four sound levels (black dots and line). (B) Proportion of facilitation trials at fifteen speaker ranks (same color dots and lines in x-axis) and four sound level ranks (different color dots and bars in y-axis) were assigned based on the response obtained under the equal-probability mode. Length of upwards arrow is proportional to the proportion of facilitation trials averaged across speakers. Slope of the northeast arrow is proportional to the proportion of facilitation trials averaged across sound levels. Dots and error bars indicate mean - standard deviation of mean. (C) Similar to (B), but for adaptation. Length of the downwards arrow is proportional to the proportion of adaptation trials averaged across speakers. Slope of the southeast arrow is proportional to the proportion of adaptation trials averaged across sound levels.

We performed the speaker rank and sound level rank analysis in 390 sessions obtained from 63 neurons. At each sound level rank, the proportion of facilitation trials tended to be larger at the lower ranked speakers (Figure 4B, fifteen same color dots in x-axis). When averaged across sound levels, the proportion of facilitation trials was only 3.4% at the speaker ranked 1st (21 sessions tested) whereas the proportion jumped to 19.1% when the test speaker was ranked 15th (41 sessions tested) (Figure 4B, slope of northeast arrow; Figure S7B, orange dots). In contrast, at each speaker rank, the proportion of facilitation trials did not vary much when the sound level rank changed (Figure 4B, four different color dots in y-axis). When averaged across speakers, the proportion of facilitation trials was 12.4% at the sound level ranked 1st (119 sessions tested) whereas the proportion rose slightly to 13.4% when the sound level was ranked 4th (61 sessions tested) (Figure 4B, length of upwards arrow; Figure S7C, orange dots).

Compared to neural facilitation, the proportion of adaptation trials tended to be larger at both higher ranked speakers (Figure 4C, fifteen same color dots in x-axis) and higher ranked sound levels (Figure 4C, four different color dots in the y-axis). When averaged across sound levels, the proportion of adaptation trials was 18.0% at the speaker ranked 1st and the proportion dropped to 8.9% when the test speaker was ranked 15th (Figure 4C, slope of southeast arrow; Figure S7B, blue dots). When averaged across speakers, the proportion of adaptation trials was 18.5% at the sound level ranked 1st and the proportion dropped to 12.3% when the sound level was ranked 4th (Figure 4C, length of downwards arrow; Figure S7C, blue dots).

These results suggest that neural facilitation is primarily dependent on sound location but not firing rate. In contrast, neural adaptation is primarily dependent on firing rate which is expected from previous adaptation literature.

### Dependence of neural facilitation on location continuity

As we have shown above, neural facilitation can be induced in the continuous presentation mode in which the probability of stimuli delivered from a target speaker is 100% (none from other speakers). In the equal-probability presentation mode, the presentation probability for each speaker is equal to 1/ 15 (number of speakers). Therefore, the continuous presentation mode provides the location continuity for sound delivery, whereas the equal-probability presentation mode does not. We further investigated in a subset of units whether the sound location continuity was necessary to induce neural facilitation by changing the presentation probability of the target speaker from 100% to 75%, 50%, 25% and 6.7% (Figure 5A, target speaker: orange square; other speakers: blue color shapes). An example neuron is shown in Figure 5B. The decrease of the target speaker’s presentation probability to 75%, 50%, and 25% resulted in increasingly weaker responses of the target speaker (100%: 20, 75%: 20, 50%: 10, and 25%: 6 spikes/sec). Across all 57 test sessions in 12 neurons, the decrease of the presentation probability of the target speaker from 100% to 75%, 50%, 25% and 6.7% resulted in weaker response of the target speaker (6.81, 3.43, 1.93, 1.89 and 0.88 spikes/sec, Figure 5C) and the reduction of the proportion of facilitation trials (44%, 23%, 16%, 21% and 14%, Figure 5D). The proportion of facilitation trials lasting for at least 5 consecutive trials showed a similar trend of decrease (Figure 5E, 25%, 12%, 5%, and 0%).

**Figure 5.**
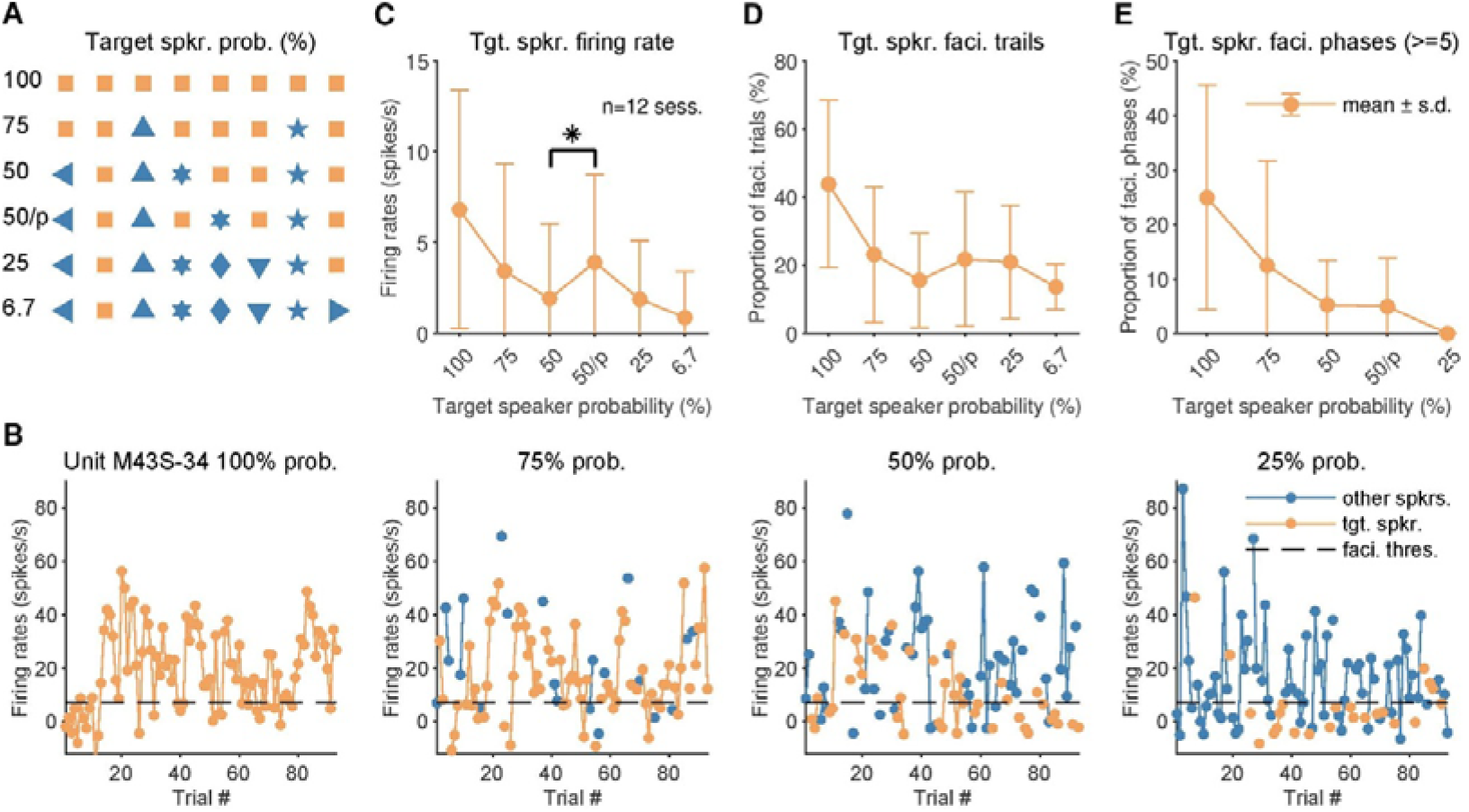
Dependence of neural facilitation on location continuity. (A) Six stimulus presentation modes were used in our studies. Continuous presentation mode equals to 100% probability presentation mode. 75%, 50% and 25% probabilistic presentation modes play sounds from all fifteen speakers in a randomly shuffled order, while giving the target speaker (orange square) a presentation probability higher than other speakers (blue left-pointing triangle, upward-pointing triangle, hexagram, diamond, downward-pointing triangle, pentagram, right-pointing triangle). For the 50% probability periodic presentation mode (50/p%), the target speaker was interleaved with other speakers. Therefore, the sequence of the target speaker was periodic instead of random. Equal-probability presentation mode equals to 6.7% probability presentation mode. (B) Firing rate of target speaker (orange dots and lines) and other speakers (blue dots and lines) for example unit M43S-34 under 100% (left), 75% (middle-left), 50% (middle-right), and 25% (right) probability presentation modes. Black dashed line indicates the facilitation threshold. (C) Firing rate of target speaker for six presentation modes. *, p<0.05, rank-sum test. Dots and error bars indicate mean ± standard deviation. (D) Proportion of facilitation trials that belong to all facilitation phases at target speaker. (E) Proportion of facilitation trials that belong to facilitation phases that lasting at least five consecutive trials at target speaker.

Previous stimulus-specific adaptation (SSA) related studies found that neurons in the auditory cortex of anesthetized rats were sensitive to statistical regularities: standard and deviant tones in random sequences both evoked larger responses than the same tones in periodic sequences (Yaron et al., 2012; Parras et al. 2017). To investigate whether neural facilitation is also sensitive to the statical regularity of the target speaker, we played two different types of sequences with 50% probability of the target speaker. In one sequence, stimuli from the target speaker and other speakers were randomly arranged, similar to the 75%, 25%, and 6.7% probability mode (Figure 5A, 50). In the other sequence, stimuli from the target speaker were interleaved with stimuli from other speakers, so that the stimuli from the target speaker were periodical (Figure 5A, 50/p). In contrast to SSA, we found that the periodic target speaker sequence evoked significantly stronger responses than the random target speaker sequence (Figure 5C, 3.91 vs. 1.93 spikes/sec, p = 0.0443, rank-sum test). The proportions of facilitation trials were similar between random and periodic target speaker sequence with 50% probability. (Figure 5D, 5E).

### Repetitive sound stimulation induced sustained membrane potential depolarization

We next asked what are cellular mechanisms underlying the neural facilitation evoked by repetitive stimuli. In a subset of experiments, we performed intracellular recordings in awake marmoset (Gao et al. 2016; Gao and Wang, 2019) to examine both membrane potential and spiking activity during both equal-probability presentation and continuous presentation modes. Figure 6A shows membrane potential traces of an example neuron. Figure 6B shows the speaker rank based on firing rate obtained under the equal-probability presentation mode in this neuron. Our previous analyses of the spiking activity measured the “facilitation phase” to characterize trials with firing rates exceeding the facilitation threshold which was defined as one standard deviation above the baseline firing rate (Figures 2A, 2D, 3D, and 6B, orange bar). Here, we measured the “depolarization phase” to characterize trials with membrane potential exceeding the depolarization threshold which was defined as one standard deviation above the baseline membrane potential (Figure 6C, blue bar). The membrane potential steadily increased after 40 trials during the continuous presentation (Figure 6D, blue line) which was accompanied by an increase in spiking activity (Figure 6D, orange line).

**Figure 6.**
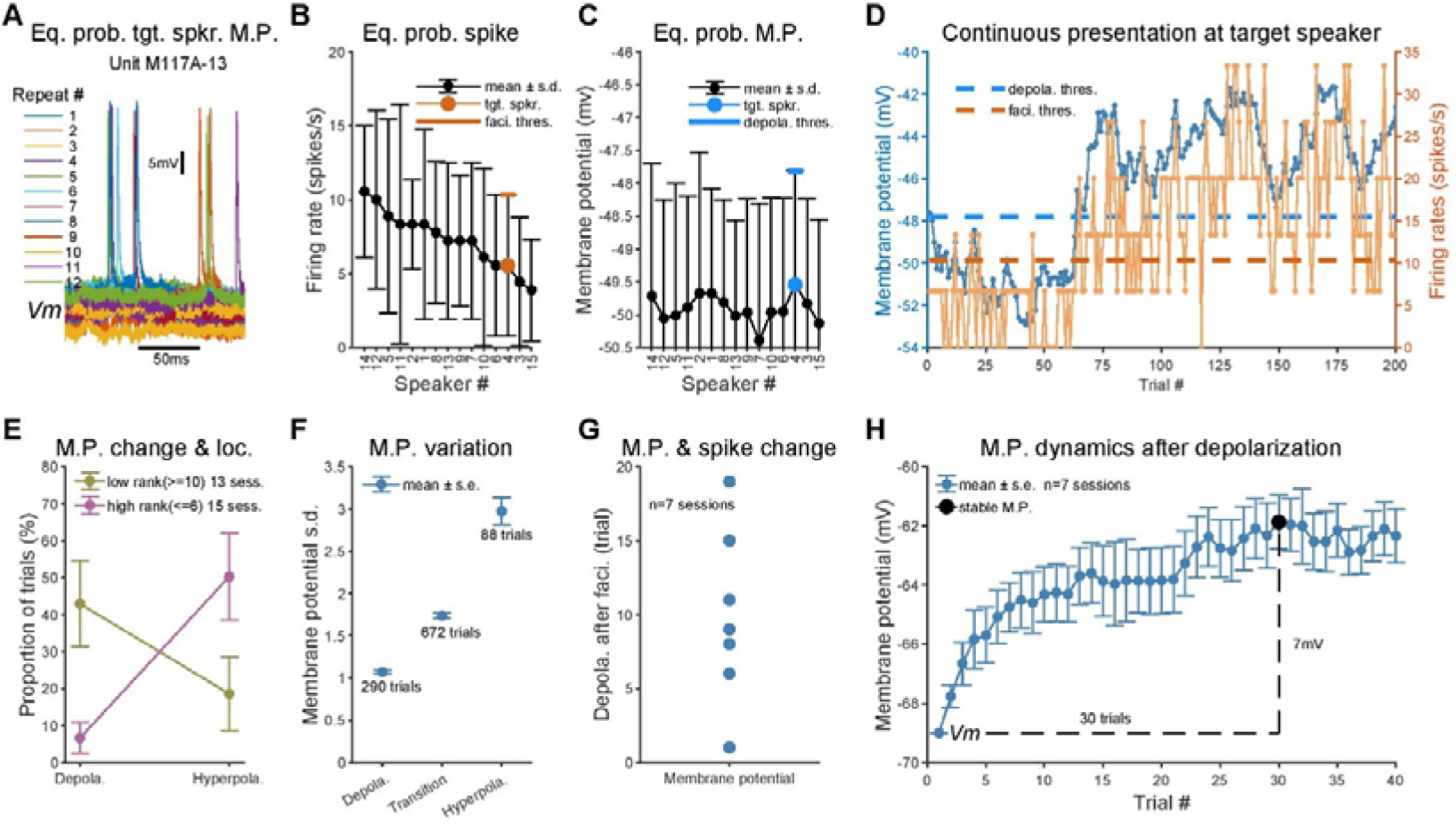
Repetitive sound stimulation induced sustained membrane potential depolarization. (A) Membrane potential traces of unit M117A-13 recorded under the equal-probability presentation mode at the target speaker location. Each color line indicates one trial. (B) Firing rate based speaker rank of same neuron obtained under the equal-probability presentation mode. Orange dot and bar indicate the target speaker and its facilitation threshold. (C) Corresponding membrane potential (spikes removed) at the same speaker rank used in (B). Blue dot and bar indicate the target speaker and its depolarization threshold. (D) Membrane potential (blue line and dots) and total firing rate (i.e., not minus the spontaneous firing rate, orange line and dots) changes during the continuous presentation mode. Threshold of facilitation (dashed orange line) and depolarization (dashed blue line) were calculated in (B) and (C), respectively. (E) Target speakers were divided into a low ranked group (10th to 15th, gold dots and bars) and a high ranked group (1st to 6th, violet dots and bars). Dots and error bars indicate mean ± standard deviation of mean. (F) The standard deviation of membrane potential for depolarization, transition and hyperpolarization trials across all sessions. (G) Number of trials needed to achieve the first membrane potential depolarization phase and the spiking facilitation phase that both lasting at least five consecutive trials. (H) Changes in membrane potential magnitude after depolarization (blue dots and line). Black dot indicates the stabilized membrane potential.

We conducted 30 sessions under the continuous presentation mode in 14 intracellularly recorded neurons at different speaker locations. The target speakers were divided into a low ranked group (10th to 15th) and a high ranked group (1st to 6th). For the low ranked group, we observed a larger proportion of depolarization trials than hyperpolarization trials (Figure 6E, gold dots, 43% vs. 19%, p = 0.1810, rank-sum test). In contrast, the opposite trend was observed for the high ranked group (Figure 6E, violet dots, 7% vs. 50%, p = 0.0044, rank-sum test). Thus, both the firing rate facilitation and membrane potential depolarization were influenced by the speaker rank. We further calculated the membrane potential variation of each trial. Trials were classified into depolarization, hyperpolarization, or transition groups based on whether they passed the depolarization threshold, hyperpolarization threshold, or neither. Depolarization trials had the lowest variation and hyperpolarization trials had the highest variation (Figure 6F, 1.07 vs. 2.97, p < 0.0001, one-way ANOVA).

For each session, we compared the difference between the number of trials needed to achieve the first membrane potential depolarization phase and the spiking facilitation phase that lasted for at least 5 consecutive trials. We found that the depolarization of membrane potential always preceded the facilitation of spiking activity (Figure 6G, median: 9 trials). Figure 6H shows changes in membrane potential magnitude after depolarization (seven sessions). The resting membrane potential was −69 mV. After 30 trials, membrane potential achieved a stable level of −62 mV which was 7 mV depolarized. We will use these three parameters in our computational models below.

### Computational models suggest two distinct neural mechanisms underlying location-specific facilitation

Our in vivo extracellular and intracellular recording data showed that both spiking and subthreshold activity of a neuron could be modulated (facilitation or adaptation of spiking activity and sustained depolarization or hyperpolarization of membrane potential) by repetitive stimuli in a location-specific way. We used two computational models to investigate potential neural mechanisms underlying these observations. In our models, synaptic inputs were panoramic (i.e., with inputs from every spatial location) based on observations from previous whole-cell recording studies in rats. Chadderton et al. (2009) found excitatory postsynaptic potential (EPSP) could be evoked from all spatial locations in all tested neurons. Kyweriga et al. (2014) found excitatory and inhibitory currents could be evoked by all interaural level difference (ILD) cues. Panoramic inputs make it possible for a cortical neuron to generate neural facilitation or sustained depolarization by amplifying weak responses at unpreferred sound locations.

A conceptual model for location-specific facilitation and adaptation in spiking activity is shown in Figure S8A. In this model, a neuron receives panoramic excitatory and inhibitory inputs but with varying strengths according to the speaker ranking as revealed by the equal probability presentation mode (upper plot). When stimuli are repetitively presented at the 15^th^-ranked speaker (right plot), stronger inhibitory inputs to the model neuron would result in larger depression than the depression of weaker excitatory inputs (Figure S8C). The overall stronger excitation than inhibition would evoke neural facilitation at this speaker location (Figure S8E). When stimuli are repetitively presented at the 1^st^-ranked speaker (left plot), stronger excitatory inputs to the model neuron would result in a larger depression than the depression of weaker inhibitory inputs. Then the overall stronger inhibition than excitation would evoke neural adaptation at this speaker location. We call this neuron model “EI-LIF model” in which excitatory and inhibitory inputs are differentially depressed (Figure 7A).

**Figure 7.**
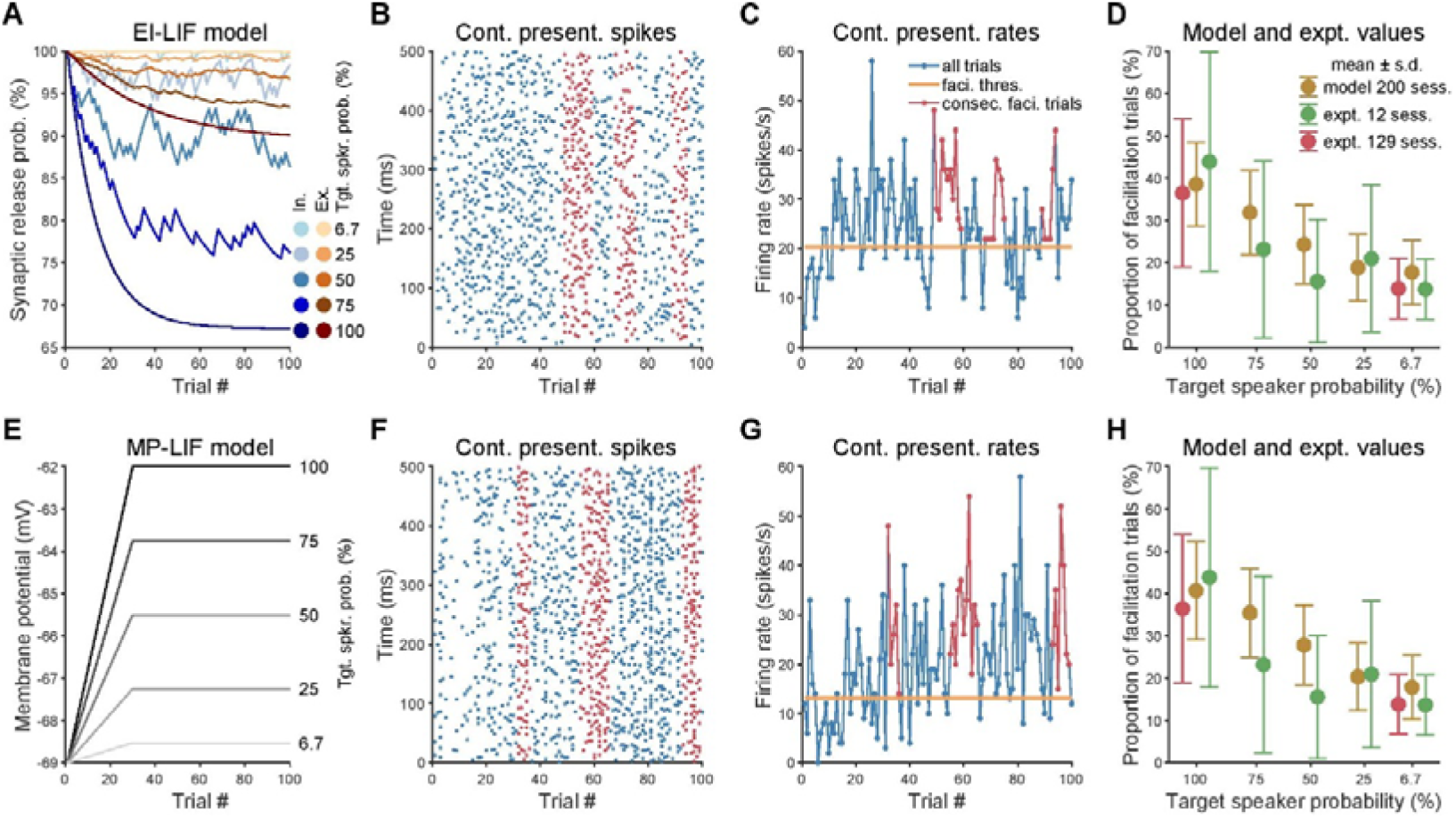
Computational models suggest two distinct neural mechanisms underlying location-specific facilitation. (A) Inhibitory (blue lines and dots) and excitatory (orange lines and dots) synaptic depression leaky integrate-and-fire model (EI-LIF model). Under the continuous presentation mode, synaptic release probability at five probability presentation modes decreased gradually. (B) Spike raster plot for one session of EI-model neuron during continuous sound presentation mode. Red dots indicate spikes belong to long facilitation phase (i.e., at least five consecutive trials with firing rates exceeding the facilitation threshold). (C) Trial-by-trial firing rate from the same session of EI-model neuron. Red dots and line indicate trails belong to the long facilitation phase. Thick orange line indicates the facilitation threshold (i.e., one standard deviation above the baseline firing rate). (D) Decrease the presentation probability of target speaker resulted in a smaller proportion of facilitation trials for 200 sessions of EI-LIF model neuron (brown dots and bars). Green dots and bars show the data from five probability presentation modes (twelve sessions, same as Figure 5D). Red dots and bars show the data from continuous and equal-probability presentation modes (129 sessions, same as Figure 3A). Dots and error bars indicate mean ± standard deviation. (E-H), Similar to (A-D) but for membrane potential depolarization leaky integrate-and-fire model (MP-LIF model).

A second conceptual model is shown in Figure S8B for membrane potential. This model is based on the idea that the brain state is modulated when stimuli switch from equal-probability presentation mode to continuous presentation mode (see Discussion). Since neural activity of the network is homeostatically regulated (Pacheco et al. 2019), sustained depolarization of weaker responses (red line) at the 15^th^ ranked speaker will be accompanied by sustained hyperpolarization of stronger responses (blue line) at the 1^st^-ranked speaker. A sustained depolarization makes it easier to reach the threshold thus evoke neural facilitation (Figures S8D, S8E). In contrast, a sustained hyperpolarization makes it harder to reach the threshold thus evokes neural adaptation. We call this neuron model “MP-LIF model”.

Figures 7B, 7C, 7F, 7G show the neural facilitation simulated with the EI-LIF model (up) and MP-LIF model (down) in the continuous presentation mode. Notice the elevated firing rates were observed in tens of trials after the start of continuous sound stimuli (Figures 7B, 7F, red dots) and epochs consisting of consecutive trials with higher firing rates than the facilitation threshold (Figures 7C, 7G, red lines). We also simulated neural adaptation in the EI-LIF model (Figure S8F) and MP-LIF model (Figure S8G). The threshold of neural facilitation and adaptation were calculated in the equal-probability presentation mode (gray area). Notice the sparse spikes (Figures S8F, S8G, blue dots) and epochs consisting of consecutive trials with firing rates lower than the adaptation threshold (Figures S8F, S8G, blue line).

We further examined whether our models could also simulate the speaker probability-dependent neural facilitation observed in Figure 5. We hypothesized that the recovery time of excitatory and inhibitory synaptic release and amplitude of sustained membrane potential depolarization were proportional to the probability of the target speaker (Figures 7A, 7E). We found that decreasing the target speaker’s presentation probability reduced the proportion of facilitation trials in the EI-LIF model (Figure 7D, gold dots) and MP-LIF model (Figure 7H, gold dots). Two models’ performances were close to the experimental results observed in Figure 5D (now shown as Figures 7D, 7H, green dots). We further compared the proportion of facilitation trials in our models with 129 experimentally tested sessions in the continuous and equal-probability presentation mode shown in Figure 3A (now shown as Figures 7D, 7H, red dots). The results were also similar at 100% and 6.7% probabilities (EI-LIF: 39% vs 36% and 18% vs 14%; MP-LIF: 41% vs 36% and 18% vs 14%). In summary, both EI-LIF and MP-LIF models recapitulated the neural facilitation and adaptation observed in our experiments and suggest potential underlying neural mechanisms. Future experimental studies can provide validations of these suggested mechanisms.

## DISCUSSION

In this study, we investigated extracellular and intracellular neural responses to repetitive sound stimulation in the auditory cortex of awake marmoset monkeys. The major finding of this study is the observation of a novel location-specific facilitation (LSF) which is dependent on sound location and stimulus presentation mode. LSF raises questions on the conventional definition of the spatial receptive field as being a static property of an auditory cortical neuron. LSF is a different phenomenon than the well-studied stimulus-specific adaptation (SSA). Computational models based on the synaptic depression or sustained depolarization mechanisms can both reproduce the LSF. The dependence of facilitation on location continuity and regularity suggests that LSF is a potential single-neuron substrate of auditory streaming.

### Implications for the spatial receptive field (SRF) of cortical neurons

The concept of a stable spatial receptive field (SRF) has been a cornerstone of our understanding of spatial tuning in the central auditory system. Auditory cortex neurons in anesthetized animals exhibit predominantly broad SRFs that typically increase in size (or width) as sound level increases (Middlebrooks and Pettigrew 1981; Mrsic-Flogel et al. 2005). In contrast, studies in awake animals have reported restricted SRFs which do not increase or show less increase in size as sound level increases (Mickey and Middlebrooks 2003; Woods et al. 2006; Zhou and Wang, 2012; Remington and Wang, 2019). It has been shown that behavior engagement could further decrease the size of SRFs and therefore sharpen spatial tuning of cortical neurons (Lee and Middlebrooks, 2011; van der Heijden et al., 2018). The finding of the present study further showed that the spatial tuning of auditory cortex neurons in awake marmosets is not static in that a non-preferred spatial location could become responsive under particular conditions. This suggests that cortical neurons can respond to spatial locations away from the center of SRF dynamically. When stimuli from other locations were inserted into the repetitively presented sound sequence from one location, neural facilitation was interrupted and even diminished (Figure 5). However, neurons still preserve their original SRF after being presented with repetitive sound stimuli (Figure S6). In contrast, after a repetitive pure tone stimulus is presented, a neuron changes its spectral receptive field by reducing responses to the specific tone frequency (Condon and Weinberger, 1991). A non-static SRF could play a role in spatial and binaural tuning plasticity that is observed in monaural deprived animals (Popescu and Polley, 2010; Keating et al., 2015).

### Comparison with stimulus-specific adaptation (SSA) and prediction

Adaptation to repetitive sound stimulation (i.e., a reduction in response to a high-probability stimulus) by auditory neurons is a commonly observed phenomenon and has been referred to as stimulus-specific adaptation (SSA) (Harpaz et al., 2021; Malmierca et al., 2014; Nelken, 2014). In previous studies that demonstrated SSA, both close field and free field sound stimulation paradigms were used in anesthetized or awake animals. In close field stimulation, the sound was delivered through a sealed speaker into the contralateral ear (Condon and Weinberger, 1991; Yaron et al., 2012; Hershenhoren et al., 2014; Nieto-Diego and Malmierca, 2016) or preferred ear (Ulanovsky et al., 2004). In free field stimulation, the sound was played from the location contralateral to (Chen et al., 2015; Kato et al., 2015; Natan et al., 2017), in front of (Natan et al., 2015) or above (Farley et al., 2010) an animal. Adaptation to repetitive sound stimulation was also observed in the current study, in particular when stimuli were delivered from preferred sound locations (i.e., higher ranked speaker locations at or near the center of SRF) (Figures 3E, 3F) or sound levels (i.e., higher ranked sound levels) (Figure 4C). However, when sound was delivered from locations away from the center of SRF, our study revealed neural facilitation to repetitive sound stimulation in the auditory cortex of awake marmosets. To the best of our knowledge, no previous studies have systematically tested repetitive sound stimulation across spatial locations.

The most striking difference between the current study and previous SSA studies is the observation of facilitation instead of adaptation to repetitive stimulation in the auditory cortex. Although the predominant response in auditory system to high probability stimuli is adaptation, unadapted and even facilitated responses have been observed in a few previous studies. Thomas et al. (2012) found that repetitive stimulation did not elicit adaptation in specialized (FM-selective) neurons of bat inferior colliculus (IC). They argued that because in echolocating bats behaviorally relevant sounds are echoes from objects, adaptation to those repetitive echolocation signals that occur with a high probability would be maladaptive during active echolocation. Parras et al. (2017) showed that most neurons on the ascending auditory pathway (IC, auditory thalamus, auditory cortex) of anesthetized rats exhibited repetition suppression. However, repetition enhancement was observed in all three areas. Lesicko et al. 2022 and Kommajosyula et al. 2021 also found repetition enhancement in the IC and auditory thalamus of awake rodents, respectively. This is consistent with our findings that repetitive stimuli evoked facilitation, and periodic target speaker sequences evoked stronger responses than random target speaker sequences (Figure 5C).

In the predictive coding model, the sensory system constantly compares external sensory inputs with internally generated predictions and updates expectations of the environment (Keller and Mrsic-Flogel, 2018). This model requires two classes of neurons: prediction error (PE) neuron and prediction (P) neuron. PE neuron signals a mismatch between predicted and actual sensory inputs, whereas the P neuron signals an expectation of sensory inputs. PE neuron has been observed in the auditory (Eliades and Wang, 2008; Keller and Hahnloser, 2009; Schneider et al., 2018), visual (Keller et al., 2012; Uran et al., 2022) and somatosensory (Ayaz et al., 2019) cortices. In contrast, the P neuron originates from other cortical areas and projects the prediction signal to the PE neuron (Schneider et al., 2014; Leinweber et al., 2017; Garner and Keller, 2022). Instead, our findings suggest that PE (i.e., repetition adaptation) and P (i.e., repetition facilitation) neurons coexist in the auditory cortex. The observation of auditory cortex inactivation reduces or blocks the repetition enhancement in the auditory midbrain and thalamus indicates prediction signals originate from P neurons in the auditory cortex (Lesicko et al., 2022; Kommajosyula et al., 2021).

Together, our finding of neural facilitation to the repetitive sound stimulation provides a complementary contextual modulation effect to the SSA and a new perspective on our current understanding of cortical responses to repetitive stimuli.

### Neural mechanisms underlying LSF

We investigated the neural mechanisms underlying LSF with two approaches. Experimentally, we directly recorded the membrane potential from neurons that exhibited LSF in awake marmosets (Figure 6). Computationally, we built a leaky integrate-and-fire (LIF) neuron model and manipulated its excitatory-inhibitory synaptic depression amplitude and recovery time (EI-LIF model) and membrane potential depolarization or hyperpolarization amplitude (MP-LIF model). Both models reproduced LSF and recapitulated the key properties of LSF (Figure 7).

Two mechanisms may account for the LSF observed in this study. One is repetitive stimulus-evoked synaptic depression. Although excitatory and inhibitory synapses are both depressed by repetitive stimuli (Galarreta and Hestrin, 1998), inhibitory synapses may show a larger amplitude of depression than excitatory synapses (Heiss et al., 2008). The imbalance between excitation and inhibition may produce LSF. Two lines of evidence support this hypothesis. First, the subthreshold activity could be evoked from all tested sound locations (Chadderton et al., 2009) and strong sound-evoked inhibition is commonly observed outside of the SRF (Zhou and Wang, 2014; Remington and Wang 2019). Therefore, the panoramic and strong inhibitory inputs are more susceptible to depression than excitatory inputs. Second, compared to the excitatory inputs, the inhibitory inputs are more sensitive to context change (Kuchibhotla et al., 2016) and show stronger depression during the forward masking (Wehr and Zador, 2005), suggesting the inhibitory inputs are more adjustable than excitatory inputs. Our EI-LIF model could reproduce the LSF based on the hypothesis that neuron has a stronger depression of their inhibitory inputs than excitatory inputs, thereby supporting a synaptic depression mechanism.

Another mechanism is salient stimulus-evoked membrane potential depolarization. A salient auditory spectrotemporal feature could attract attention automatically (Kayser et al., 2005; Huang and Elhilali, 2020). A wideband noise used in this study was not salient when it was presented randomly from different locations to characterize SRF. However, a wideband noise could become salient when it was repetitively presented from one location while sounds at all other locations disappeared. This auditory spatial pop-out hypothesis is similar to visual saliency where a visual item in sharp contrast with its neighboring items in a simple feature, such as color or orientation, automatically captures attention (Yan et al., 2018). If a location-specific wideband noise is salient, a more salient sound feature at the spectrotemporal domain, e.g., amplitude-modulated wideband noise and vocalization, indeed evoke a larger proportion of facilitation trials than the less salient unmodulated wideband noise (Figure 3C). In humans, salient auditory stimuli dilate the pupil (Wang et al., 2014). Pupil dilation is closely correlated with membrane potential depolarization and a decrease in membrane potential variation (McGinley et al., 2015). Interestingly, we observed similar changes in membrane potential when neurons exhibit LSF (Figure 6), suggesting that the ongoing repetitive stimuli are salient to the animals. Importantly, our MP-LIF model reproduced LSF by incorporating parameters obtained from intracellular recordings. Together, our intracellular recording data and MP-LIF model simulation support a salient stimulus-evoked sustained depolarization mechanism underlying the LSF.

It is not clear whether top-down attention plays a role in LSF. It has been shown that task engagement could modulate the SRFs (Lee and Middlebrooks, 2011). In the current study, marmosets passively listened to sound stimuli, though they might have chosen to pay attention to repeated stimulation from a particular location in the continuous stimulation mode. However, we did not observe an increase in spontaneous activity when neural facilitation was observed (Figure S6F), whereas attention tends to increase spontaneous activity (Luck et al., 1997; Reynolds et al., 2000). Furthermore, we found no preference of different cell types in exhibiting the LSF (Figure S3D), whereas top-down attention has stronger modulation over putative inhibitory neurons (Mitchell et al., 2007).

### Candidate neural substrate for auditory streaming

In a natural environment like at a cocktail party, sounds are often simultaneously and continuously generated by multiple sound sources (Cherry, 1953). One major challenge for a listener is forming auditory streaming (McDermott, 2009). Streaming requires acoustic cues such as frequency, temporal regularity, and sound location (Shamma and Micheyl, 2010). Over the past decade, a rapidly increasing number of studies have investigated the effect of temporal regularity or repetition for streaming (Bendixen et al., 2010; Andreou et al., 2011). Regular stimulation induces stronger responses than random stimulation when measured with magneto-electro encephalography (M/EEG) and functional MRI (fMRI) in humans (Barascuda et al., 2016; Southwell et al., 2017). The findings that repetitive sound stimulation evoked LSF and regular stimulation evoked a stronger response than random stimulation provide a candidate single-neuron correlate of this perceptual phenomenon. Computational modeling suggests that the change of synaptic efficacy could result in sustained responses to regular stimulation (Auksztulewicz et al., 2017). Interestingly, changing the excitatory and inhibitory synaptic efficacy in our EI-LIF model also generated the LSF. Those two models further suggest that sustained response to regular stimulation and LSF share a similar neural mechanism. Repetition causes the target to pop out from the background and is robust to inattention (McDermott et al., 2011; Masutomi et al., 2016; Mehta et al., 2021). Based on the same pop-out hypothesis, our MP-LIF model could reproduce the LSF. Those similarities suggest that our EI-LIF and MP-LIF models provide a theoretical foundation for both LSF observed in marmosets and enhanced response to regular over random stimulation observed in humans. Together, our findings and models provide valuable new insights into the neural mechanisms of auditory streaming.

## METHODS

### Animal preparation and experimental setup

Data were collected from five hemispheres of four monkeys (Monkey 1: left, Monkey 2: left and right, Monkey 3: right, Monkey 4: left). All experimental procedures were approved by the Johns Hopkins University Animal Use and Care Committee. These procedures were identical to those described in previous publications from our laboratory (Lu et al. 2001). A typical recording session lasted 3-4 h, during which an animal sat quietly in a specially adapted primate chair with its head immobilized. Throughout the entire recording session, the animal was closely monitored via a video camera by the researcher. The eye position was not controlled, but when the animal closed its eyes for a prolonged period, the experimenter ensured the animal opened its eyes before the next stimulus set was presented.

Experiments were conducted in a double-walled sound-proof chamber (Industrial-Acoustics) with the internal walls and ceiling lined with three-inch acoustic absorption foam (Sonex). Fifteen free-field loudspeakers were placed on a semi-spherical surface centered around the animal’s head and above the horizontal plane. The speaker setup was similar to our previous studies (Zhou and Wang, 2012), but with speakers covering the rear sphere. Eight speakers were evenly positioned at 0° elevation, five speakers were evenly spaced at +45° elevation in the frontal hemifield, one speaker was located at +45° elevation in the rear midline, and one speaker was located directly above the animal.

### Extracellular and intracellular recordings

Extracellular recording procedures were identical to those described in our previous publications. A sterile single tungsten microelectrode (A-M Systems) was held by a micro-manipulator (Narishige) and inserted nearly perpendicularly into the auditory cortex through a small opening on the skull (1.0–1.1mm in diameter) and advanced by a hydraulic micro-drive (David Kopf Instruments). The tip and impedance of the electrode was examined before each recording session (2-5MΩ impedance). Spikes were detected by a template-based spike sorter (MSD, Alpha Omega Engineering) and continuously monitored by the experimenter while data recordings progressed. The raw voltage signal was also recorded. Intracellular recording procedures were identical to those described in our previous publications (Gao et al. 2016; Gao and Wang, 2019). The recordings were made in the auditory cortex through the intact dura using a concentric recording electrode and guide tube assembly. The sharp recording pipette was made of quartz glass. The guide tube was made of borosilicate glass. The sharp recording pipette was pulled by a laser puller (P-2000, Sutter Instrument), and the guide tube was pulled by a conventional puller (P-97, Sutter Instrument). The electrode assembly was advanced perpendicularly relative to the cortical surface with a motorized manipulator (DMA1510, Narishige). The electrical signals were amplified (Axoclamp 2B, Molecular Devices), digitized (RX6, Tucker-Davis Technologies), and saved using custom programs (MATLAB, Mathworks).

### Acoustic stimuli

Four different stimulus presentation designs were used. 1) Continuous (100%): the same stimulus was repeatedly delivered from a fixed speaker location over many trials. 2) Unequal-probability and random (75%, 50%, and 25%): stimulus was delivered from multiple speaker locations in a randomly shuffled order, but target speaker has a higher probability than others. 3) Unequal-probability and periodic (50/p%): stimulus delivered from the target speaker was interleaved with stimulus delivered from other speakers, so that the stimulus delivered from the target speaker was periodical. 4) Equal probability (6.7%): stimulus was delivered from multiple speaker locations in a randomly shuffled order and all speakers shared the same occurrence probability. For each neuron, if allowed by the experiment conditions, the equal-probability presentation mode was tested at multiple separate sessions at different time points and other three presentation modes were tested between the equal-probability presentation mode.

Stimuli were generated digitally in MATLAB at a sampling rate of 97.7kHz using custom software, converted to analog signals (RX6, Tucker-Davies Technologies), attenuated (PA5, Tucker-Davies Technologies), power amplified (Crown Audio), and played from specified loudspeaker. The sound tokens used included unfrozen wide-band noise, frozen wide-band noise, amplitude-modulated wide-band noise and species-specific vocalizations. Sessions collected under continuous unfrozen wide-band noise stimuli were used only in Figure S4H. Fixed inter-stimulus intervals (ISI) were used in four presentation modes. A variety of random ISI were used only in continuous presentation mode. The shortest ISI was 500ms, and the longest ISI was 5200ms. Rate-level function was used to find the best sound level of tested neurons. Most neurons were tested using best sound level, except sound level rank experiments where four different sound levels were used. The same sound level was used when comparing different presentation modes.

### Characterization of spatial receptive fields

Total firing rates were calculated over a time window beginning 15ms after stimulus onset and 50ms after stimulus offset. Total firing rates subtracted by the spontaneous rate was the firing rate. SRF characterization was identical to our previous studies (Remington and Wang, 2019). The threshold was the half maximal firing rates. Tuning selectivity was defined as the number of areas that have higher firing rates than the threshold divided by the total number of areas. Direction selectivity was defined as the product of every area, unit vector and firing rate divided by the product of every area and firing rate. If a neuron only responded to contralateral and ipsilateral 90° at horizontal plane and have equal firing rates, then the direction selectivity was zero. In the plotted SRF, the location of white color dot indicated the preferred sound location, the dot diameter was proportional to the direction selectivity, the black thick line was the half-maximum threshold of SRF, area encircled by the threshold was the reciprocal of tuning selectivity.

### Identification of cortical areas, layers and cell types

We used the best frequency (BF) of neurons to identify the subregions of auditory cortex. For the neurons significantly responding to at least one tone stimulus played at the front speaker, we specified the frequency of the tone stimulus that evoked the maximum response rate as the neuron’s BF. Marmoset auditory cortex is situated largely ventral to the lateral sulcus and exhibits a topographical frequency gradient along the rostral–caudal axis. The boundary between primary auditory cortex (A1) and the caudal area (caudal-medial and caudal-lateral belt) can be identified by an abrupt decrease of BF at the high-frequency (caudal) border of A1. A1 was further divided into the low-frequency (≤8KHz) rostral A1 and high-frequency (>8KHz) caudal A1 along the rostral-caudal axis. First spike depth was the absolute depth where the first spike was detected from this electrode, and was depended on thickness of dura, variations in granulation tissue, proximity to the curvature of the sulcus, and orthogonality of the electrode penetration to the cortical surface. We used the trough to peak duration of spike waveforms to identify putative excitatory and inhibitory neurons (Mitchell et al., 2007). Before analyzing the spike duration, we calculated the signal to noise ratio of spike waveforms (Sabyasachi and Wang, 2012), which was defined as the action potential peak to peak height divided by the standard deviation of the background noise over 1ms preceding all spikes (20 × log10(AP_peak–peak_/Noise_SD_)).

### The proportion of facilitation, adaptation, depolarization and hyperpolarization trials

The facilitation phase was defined as the trials whose firing rates were at least one standard deviation above the mean firing rate evoked under the equal-probability presentation mode. The sum of facilitation trials divided by the total trial number in each session was defined as the proportion of facilitation trials. The adaptation phase was defined as the trials whose firing rates were at least one standard deviation below the mean firing rate evoked under the equal-probability presentation mode. To analyze the membrane potential change, spikes were removed from voltage signal with a time window of 3ms centered around the spike peak. The proportion of depolarization and hyperpolarization trials were calculated similar to the proportion of facilitation and adaptation trials.

### Speaker rank and sound level rank

Speaker tested in each continuous presentation mode was assigned a rank number based on its firing rate obtained under the equal-probability presentation mode. Speaker ranked 1st and 15th had the highest and lowest firing rate among all 15 speakers tested, respectively. When the SRF has a single peak, speaker ranked 1st and 15th usually had the closest and farthest distance to the preferred direction, respectively. When the SRF has multiple peaks, low ranked speaker may next to the preferred direction occasionally. We used speaker firing rate rank instead of distance rank for two reasons: one, the SRF of some neurons was quite dispersive, thus it was inaccurate to compute the distance between the target speaker and the SRF center; and two, the SRF center was usually determined by several high ranked speakers. The contribution of low ranked speakers was not considered when using the distance rank. For sound level rank, we measured neurons’ responses to 200ms wide-band noise played at four sound levels under the equal-probability presentation mode. These four sound levels were a series of fixed values with an interval of 20dB.

### Computational models of location-specific facilitation

Computational models that recaptured the location-specific facilitation (LSF) phenomenon were based on two neural mechanisms (Figures S8A-S8D): excitatory and inhibitory synaptic depression (EI-LIF model, see equations 4 and 5) and membrane potential depolarization and hyperpolarization (MP-LIF model, see equation 6 and 7). EI-LIF model dynamically changed the depression amplitude of synaptic vesicles release and exponential recovery time constant. MP-LIF model dynamically changed the resting potential and spiking threshold. In the LSF, the parameters were modulated by the probability of presentation speaker, e.g., 100% for continuous presentation mode and 6.7% for equal-probability presentation mode.

The membrane potential *V*_*t*+1_ of a leaky integrate-and-fire (LIF) neuron at time step Δ*t* was:

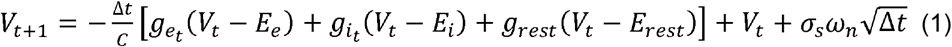

*g_e_t__* and *g_i_t__* was the excitatory and inhibitory synaptic conductance (see equations 2 and 3). *C, E_e_, E_i_* and *g_rest_* was the membrane capacitance, excitatory reversal potential, inhibitory reversal potential, and leak conductance. Those values were obtained from the *in vivo* whole-cell recording in the auditory cortex of anesthetized rats (Wehr and Zador, 2003). Gaussian noise *σ_s_ω_n_* was added to generate the spontaneous firing (Lee et al., 2020). Action potential was evoked when the *V*_*t*+1_ reached the spike threshold *V_spike_*. *V*_*t*+1_ was reset to *E_rest_* after the action potential. *E_rest_* was obtained from our intracellular studies (Figure 6H). It was fixed in the EI-LIF model but was dynamically modulated in MP-LIF model. *V_spike_* is the sum of threshold above resting potential *V_th_* and *E_rest_*. It was fixed in the EI-LIF model but was modulated in MP-LIF model.

Excitatory conductance *g_e_t__* and inhibitory conductance *g_i_t__* were:

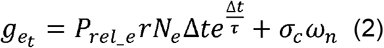

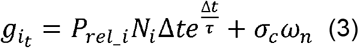

*P_rel_e_* and *P_rel_i_* were the excitatory and inhibitory synaptic release probability, respectively (see equations 4 and 5). *P_rel_e_* and *P_rel_i_* were modulated in EI-LIF model but fixed to one in MP-LIF model. The inhibitory and excitatory inputs have the same strength and occurred simultaneously, so the inhibitory to excitatory ratio *r* equal to one and the inhibitory to excitatory delay *d* (not shown in the equation) equal to zero. The number of excitatory inputs *N_e_*, inhibitory input *N_i_* and time constant *τ* were fixed and conduction noise *σ_c_ω_n_* was added to generate the spontaneous firing (Wehr and Zador, 2003).

For the EI-LIF model, in each trial *T*, the excitatory and inhibitory synaptic release probability *P*_*rel_e*_*t*+1__ and *P*_*rel_i*_*t*+1__ were:

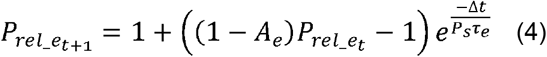

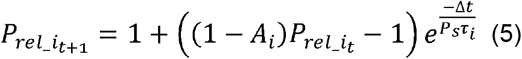

*A_e_* and *A_i_* was the excitatory and inhibitory synaptic depression amplitude, respectively. *A_i_* was larger than *A_e_* because we observed facilitation instead of adaptation in the continuous presentation mode. In addition, Heiss et al., 2008 found that inhibition adapts more than excitation when repetitively stimulating the whisker. *A_e_* and *A_i_* were both fixed in the LSF. Relatively small depression amplitude was chosen due to the slow facilitation processes observed in the recording data.

*τ_e_* and *τ_i_* was the excitatory and inhibitory recovery time constant, respectively. *τ_e_* was longer than *τ_i_*, because numerous studies found that inhibitory synapses have a quick recovery than excitatory synapses (Galarreta and Hestrin, 1998; Varela et al., 1999). In the different probability presentation mode, facilitation percent and firing rate decreased when the probability of target speaker decreased. Since the time constant was stimulus frequency dependent (Galarreta and Hestrin, 1998), therefore the *τ_e_* and *τ_i_* were scaled by the presentation probability *P_s_* which resulted in higher synaptic release probability, i.e., less adaptation, when the presentation probability was low. Time constant can across multiple time scales from hundreds of milliseconds to tens of seconds (Varela et al., 1997; Ulanovsky et al., 2004). Since 44 trials were required to reach the first long facilitation phase, *τ_e_* and *τ_i_* were chosen so that the probability of release was stable after 40 trials.

Each session is composed of randomly presented trials *T_r_* for computing the facilitation and adaptation threshold and continuously presented trials *T_c_* for computing the facilitation percent, adaptation percent, and firing rate. The median firing rate in the 100% probability presentation mode was 12 spikes per second. Therefore, the number of stimulus count *N_sc_* was chosen so that the average firing rate in the EI-LIF model could match the firing rate in the recording data. Notice that *N_sc_* was Poisson distributed and not every stimulus input could evoke a spike output in the LIF neuron. We run EI-LIF model for two hundred sessions for every presentation probability.

For the MP-LIF model, in each trial *T*, the dynamic resting membrane potential *E_rest_MP_* and spike threshold *V_spike_* were:

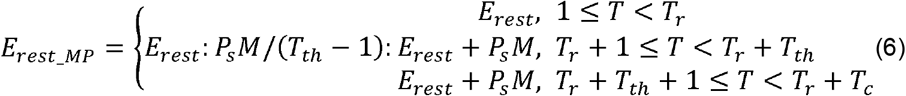

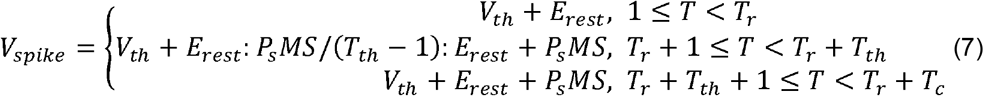

The depolarization value *M* and the number of trials to reach the stabilized depolarization value *T_th_* were obtained from our intracellular recordings. Since lower presentation probability evoked less neural facilitation, therefore *M* was scaled by the presentation probability *P_s_*. Spike threshold *V_spike_* was further modulated by the spike threshold scale S. Almost no spike was evoked when S equal to one but excessive spikes were evoked when S equal to zero. Therefore, S and stimulus count *N_sc_* were chosen so that the average firing rate matched the recording data. We also run MP-LIF model for two hundred sessions at every presentation probability. Table S1 listed the names of parameters and corresponding values used in EI-LIF and MP-LIF model neurons

## DATA AND CODE AVAILABILITY

Source data for generating Figure 1 to Figure 7 is publicly available (https://github.com/ccg1988/LSF_Figures). Codes (MATLAB R2022a) for generating Figure 1 to Figure 7 and computational models is publicly available (https://github.com/ccg1988/LSF_Figures; https://github.com/ccg1988/LSF_Model). Codes (MATLAB R2012a) for controlling the TDT sound presentation and data acquisition system are freely available upon request from the corresponding author.

## ACKNOWLEDGMENTS

This research was supported by National Institute of Health grants DC003180 and DC005808 to X.W. We thank A. Pistorio, and J. Estes for assistance with animal care, X. Song, X. Liu, and H. Mandal for their comments on the earlier versions of the manuscript.

## AUTHOR CONTRIBUTIONS

Conceptualization, S.X., C.C. and X.W.; investigation, S.X., Y.W., and C.C.; visualization, C.C.; funding acquisition, X.W.; project administration, X.W.; supervision, X.W.; data analysis, S.X. and C.C.; modeling, C.C.; writing – original draft, C.C.; writing – review & editing, C.C. and X.W.

---

**Figure S1.**
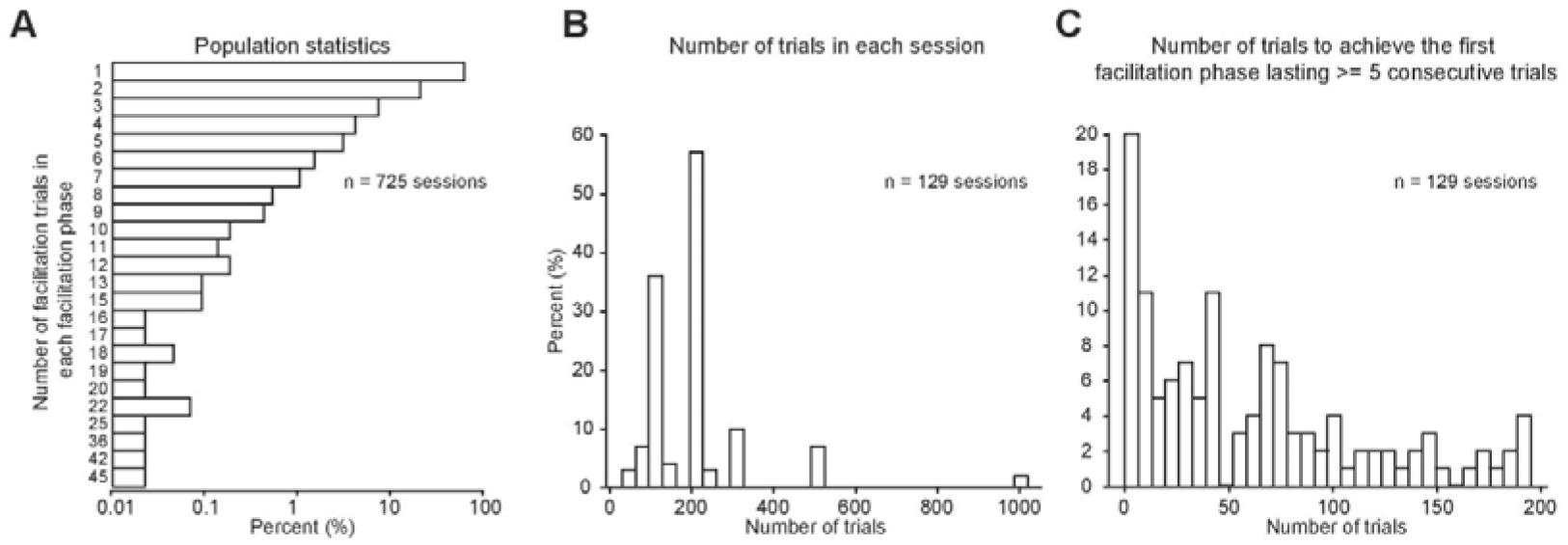
Population statistics. (A) Logarithmic histogram of the number of facilitated trials in each facilitation phase. We chose the sessions that exhibited facilitation phases lasting at least five consecutive trials as the threshold. Fifty-one neurons passed the threshold. (B) Histogram of the proportion of tested trials in each session (fifty-one neurons). (C) Histogram of the proportion of trials that was required to achieve the first facilitation phase lasting at least five consecutive trials (fifty-one neurons).

**Figure S2.**
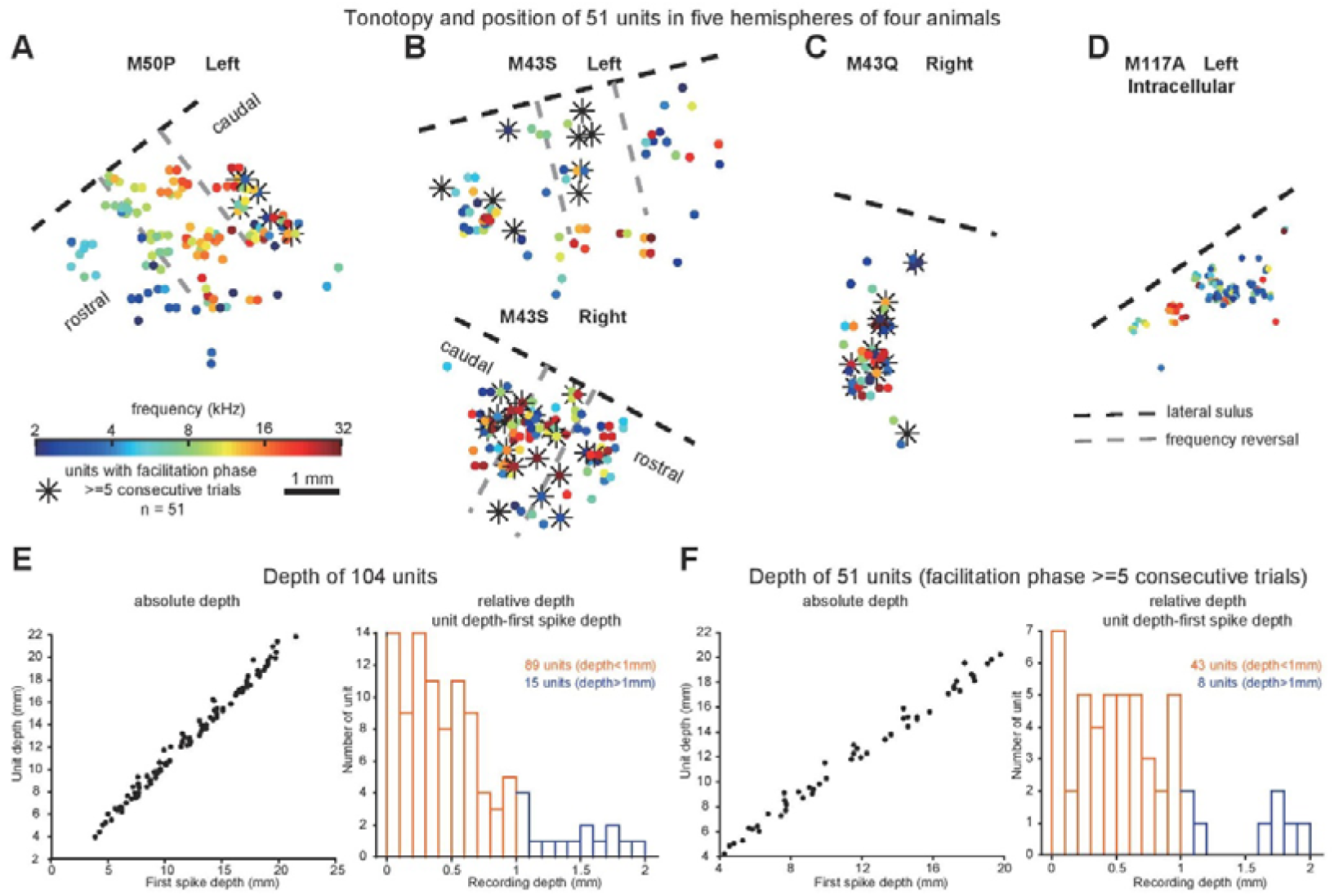
Facilitation neurons across cortical areas and layers. (A-D) Best frequency (BF) distribution on the cortical surface from five hemispheres of four monkeys. The BF of recorded neurons (color dots) was examined based on their responses to pure tones stimuli ranging from 2kHz (blue) to 32kHz (red). Fifty-one neurons that exhibited facilitation phases lasting at least five consecutive trials were labeled with asterisks. The black dashed line marked the lateral sulcus line approximated by the bone suture in the temporal lobe. The auditory cortex was divided (gray dashed line) into the rostral primary auditory cortex (A1), caudal A1, and caudal area (caudal lateral and caudal medial) along the rostral-caudal axis based on the BF distribution change. (A) Tonotopy from left hemisphere of Monkey 1. Six facilitated neurons were located in caudal area. (B) Ten facilitated neurons (two overlapped) located in A1 of left hemisphere (top) and twenty-two facilitated neurons (two overlapped) located in A1 and caudal area of right hemisphere (bottom) of Monkey 2. (C) Thirteen facilitated neurons (two overlapped) from right hemisphere of Monkey 3. (D) Fourteen neurons were recorded using the intracellular recording method in the left hemisphere of Monkey 4. (E) Depth information for all the 104 neurons. Left, first spike depth (x-axis) is the absolute depth where the first spike is detected, unit depth (y-axis) is the absolute depth where the neuron is recorded currently. Right, recording depth equals to the relative depth which is calculated as the unit depth minus first spike depth. Eighty-nine neurons came from supragranular layers (recording depth less than 1mm, orange bars). (F) Same to (E) but for fifty-one neurons that exhibited facilitation phases lasting at least five consecutive trials. Forty-three neurons came from supragranular layers (orange bars).

**Figure S3.**
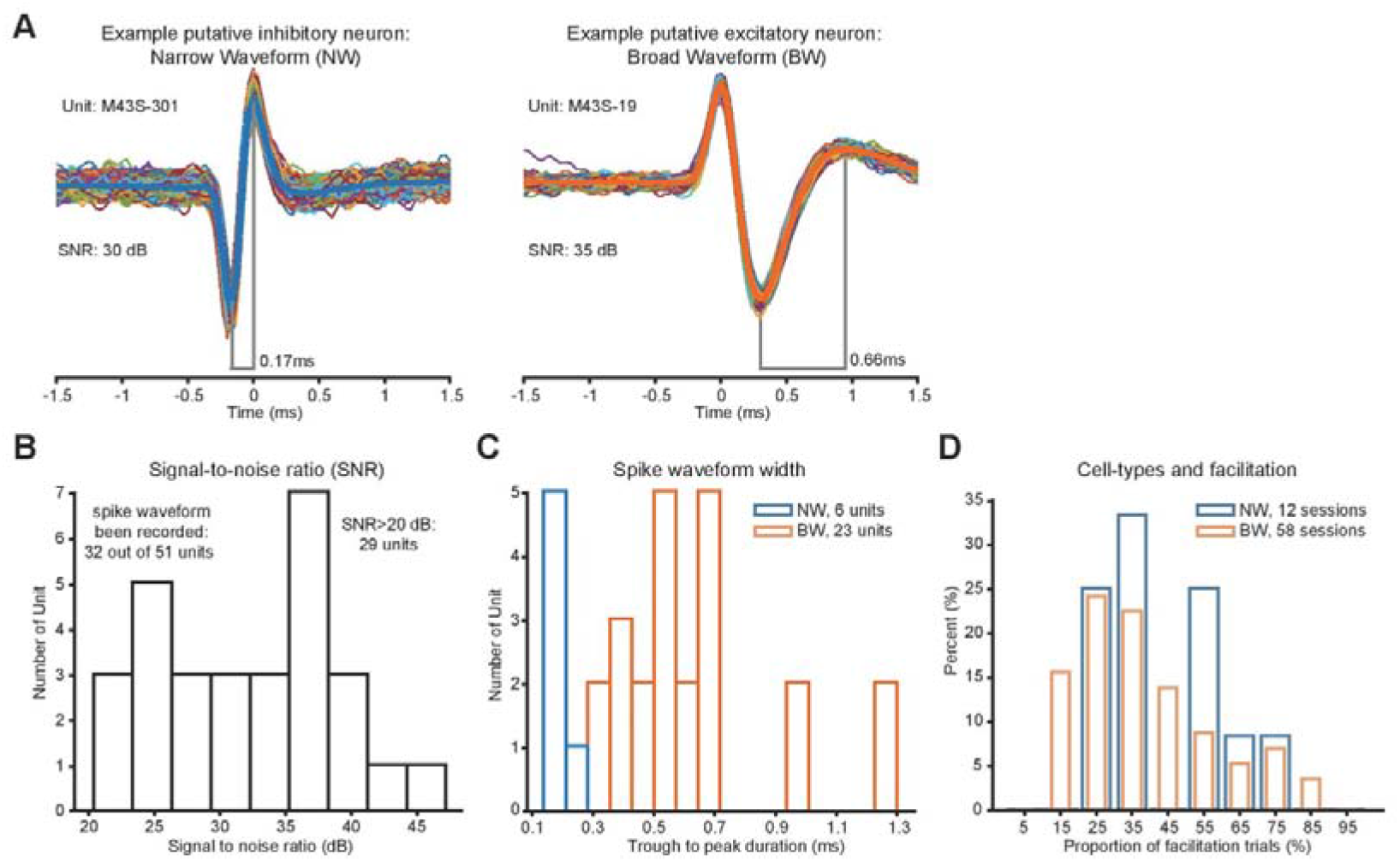
Facilitation in putative excitatory and inhibitory neurons. (A) Example putative inhibitory narrow-waveform (NW, left) and putative excitatory broad-waveform (BW, right) neurons. (B) Among the fifty-one neurons with facilitation phase lasting at least five consecutive trials, the waveform of thirty-two neurons has been recorded. Twenty-nine neurons have a signal-to-noise ratio that was larger than 20dB. (C) Among the twenty-nine neurons, we used 0.3ms trough to peak duration as the boundary for classifying putative inhibitory (blue bars) and excitatory (orange bars) neurons. 21% of neurons were classified as putative inhibitory neurons. (D) Histogram of the proportion of facilitation trials between putative inhibitory (blue bars) and excitatory (orange bars) neurons.

**Figure S4.**
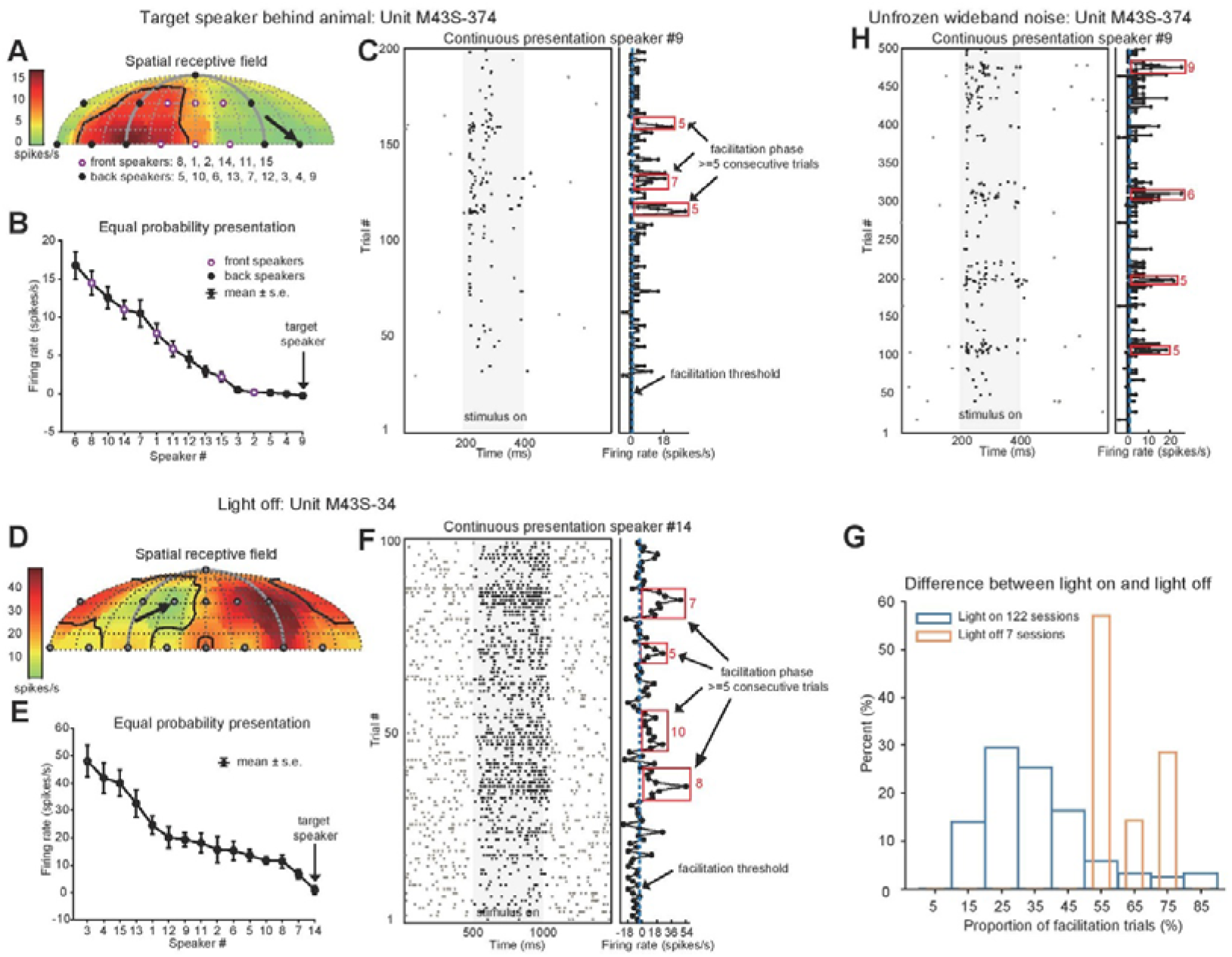
Facilitation occurred in the back, in the darkness, in using unfrozen wideband noises. **(A)** Spatial receptive field of example unit M43S-374 obtained under equal-probability presentation mode. Black arrow indicates the spatial location of continuously presented stimuli. Six orange circles indicate front speakers, and nine black dots indicate back speakers. **(B)** Firing rate versus speak number of same example neuron under the equal-probability presentation mode. **(C)** Left, spike raster of same example neuron tested at speaker #9 under continuous sound presentation mode. Gray shaded area indicates the sound presentation period. Right, trial-by-trial firing rate. Dashed blue line indicates the facilitation threshold. Red squares include trials belonging to the long facilitation phase (i.e., at least five consecutive trials with firing rates exceeding the facilitation threshold). **(D-F)** Similar to **(A-C)**, but the example unit M43S-34 was tested under the light-off condition. **(G)** Histogram of the proportion of facilitation trials under the continuous presentation mode for light on (blue bars) and off (orange bars) sessions. Among the seven light-off sessions, five sessions were tested when light was turned off, and two sessions were tested when both eyes were blocked by an acoustic drape. **(H)** Unfrozen wideband noise stimuli were tested for example unit M43S-374. Four sessions tested with unfrozen wideband noise stimuli were not included in 129 sessions mentioned above.

**Figure S5.**
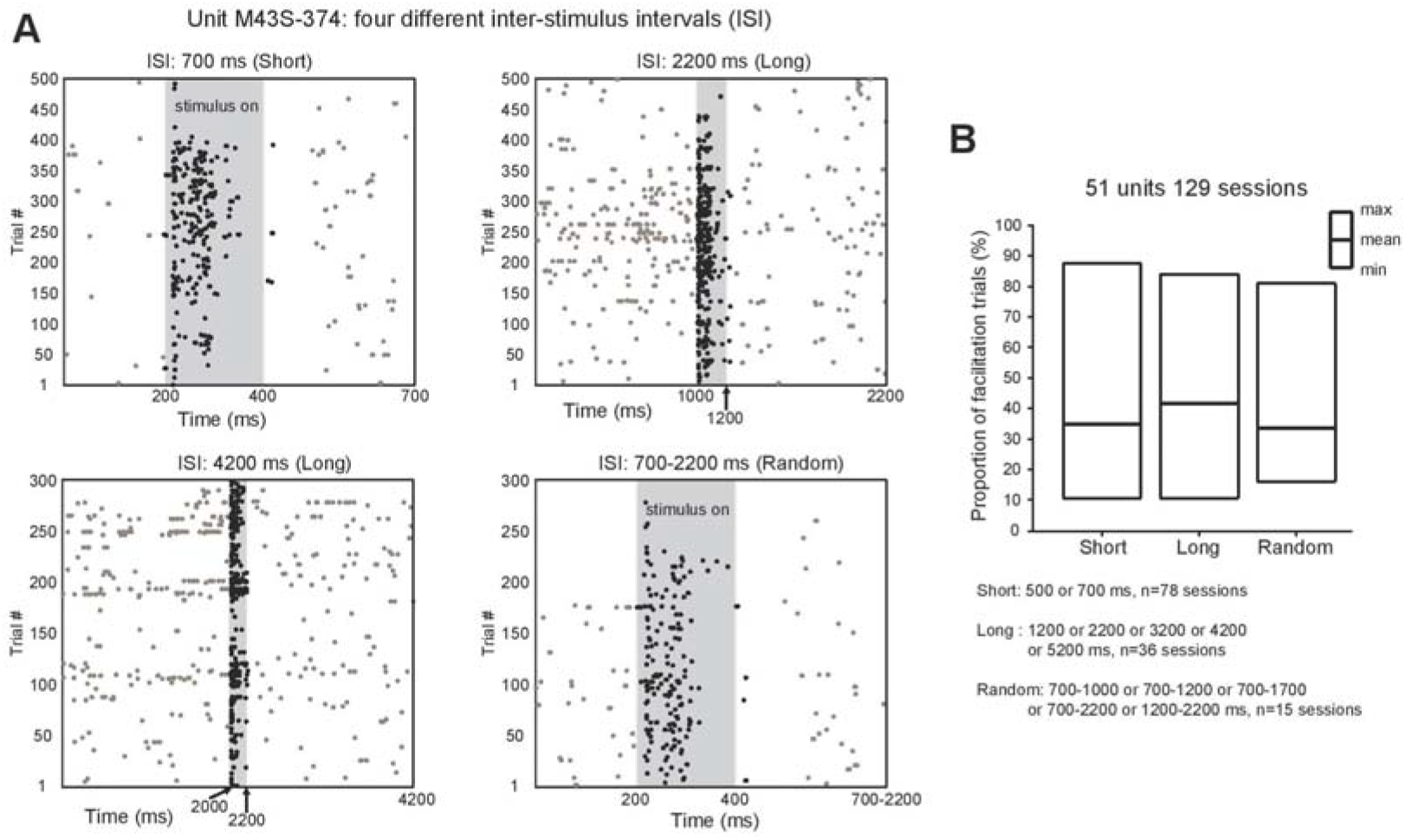
Facilitation occurred using different ISIs. (A) Example unit M43S-374 showed neural facilitation at fixed length 700ms ISI (top left), 2200ms ISI (top right), 4200ms ISI (bottom left), and random length 700-2200ms ISI (bottom right). Gray shaded area indicates the sound presentation period. (B) Across the 129 sessions with facilitation phase lasting at least five consecutive trials, ISIs were classified into three groups: short (500 and 700ms, 78 sessions), long (1200, 2200, 3200, 4200 and 5200ms, 36 sessions) and random (700-1000 or 700-1200 or 700-1700 or 700-2200 or 1200-2200ms, 15 sessions).

**Figure S6.**
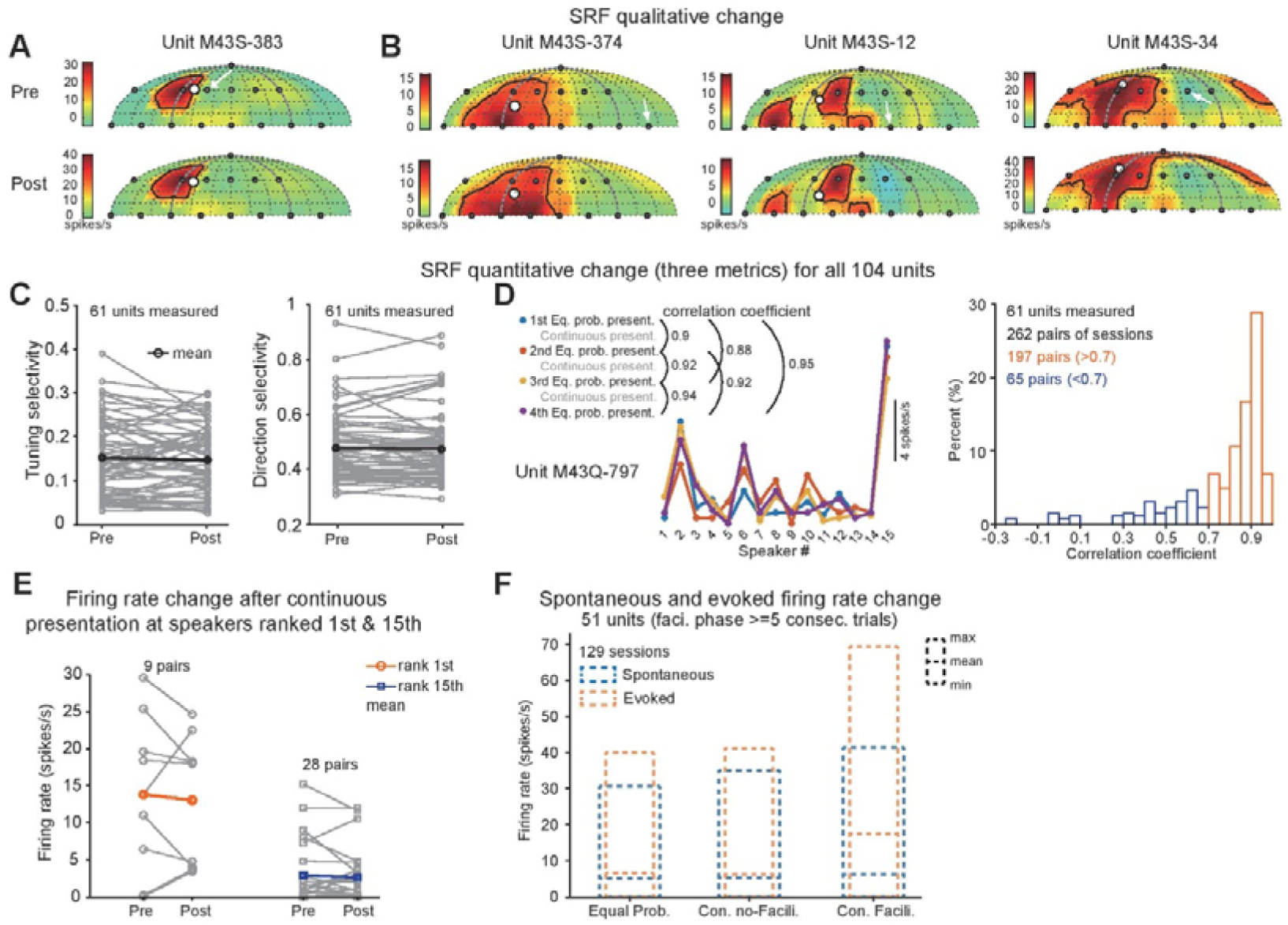
Neural facilitation did not alter SRF. (A) Spatial receptive field under the equal-probability presentation mode before (up) and after (down) the presentation of continuous sound stimuli (same neuron in Figure 1C). White arrow indicates the speaker location tested under continuous presentation mode. Area within the enclosed black line is proportional to the reciprocal of tuning selectivity, and the size of white dot is proportional to the direction selectivity. (B) Similar to (A) but for all other three example neurons. (C) Tuning selectivity (left) and direction selectivity (right) before and after the continuous presentation mode. (D) Left, the pairwise correlation coefficient between each pair of responses to fifteen speaker locations in the example unit M43Q-797. Color dots and lines indicate averaged firing rate under equal-probability presentation modes. Right, histogram of the correlation coefficient before and after the continuous presentation mode. Orange bars indicate paired sessions with a correlation coefficient greater larger than 0.7. (E) Total firing rates (i.e., without minus the spontaneous firing rate) changes for the highest (1st, orange circles and line) and lowest (15th, blue circles and line) ranked target speakers (rank 1st: p = 0.7962, rank 15th: p = 0.9738, rank-sum test). Color dots indicate the mean value. (F) Spontaneous (blue dashed boxes) and evoked (orange dashed boxes) total firing rates of equal-probability presentation mode, non-facilitation, and facilitation phases of the continuous presentation mode. For equal-probability presentation mode, only trials using the target speaker were included.

**Figure S7.**
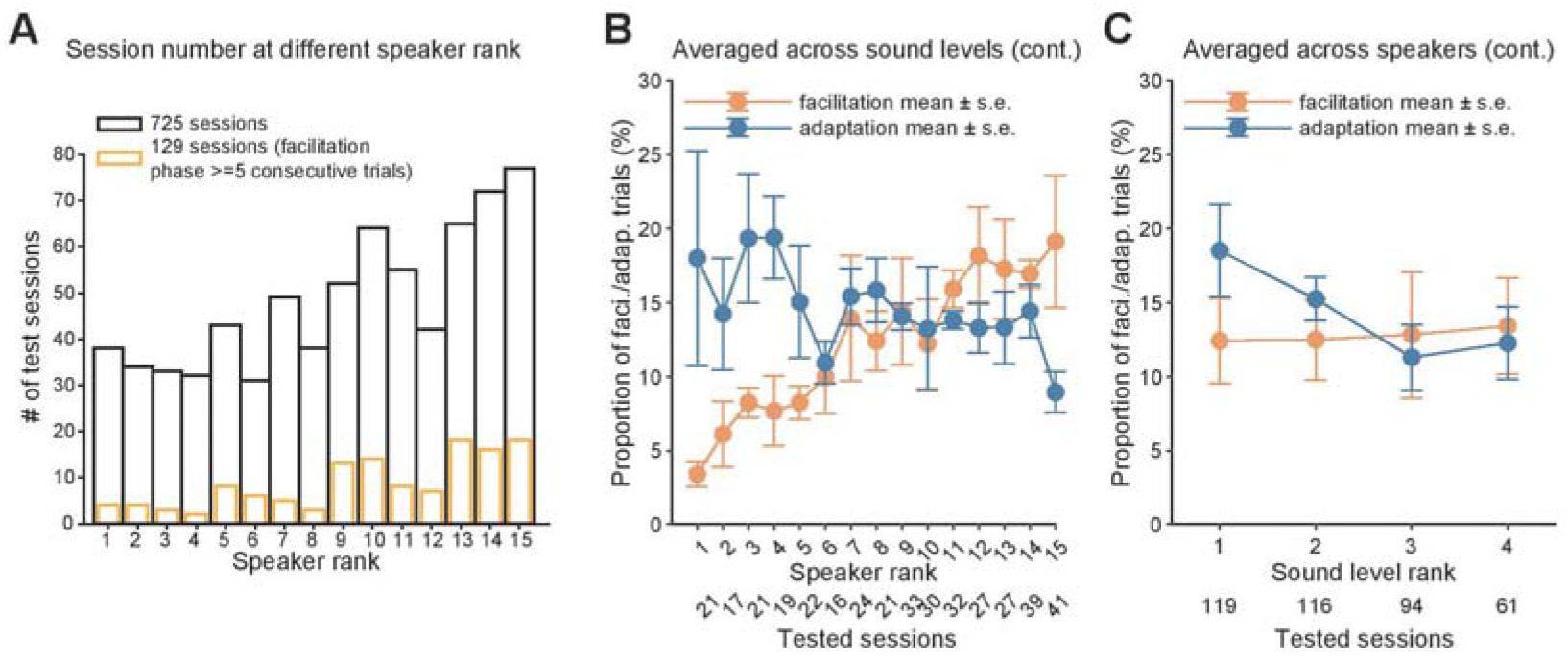
Proportion of facilitation and adaptation trials averaged across sound levels and speakers. (A) Number of all tested sessions (black bars) and sessions with facilitation phases lasting at least five consecutive trials (yellow bars) at different speaker rank. (B) Proportion of facilitation (orange dots and bars) and adaptation (blue dots and bars) trials averaged across sound levels under the continuous presentation mode. Dots and error bars indicate mean ± standard deviation of mean. (C) Proportion of facilitation (orange dots and bars) and adaptation (blue dots and bars) trials averaged across speakers under the continuous presentation mode.

**Figure S8.**
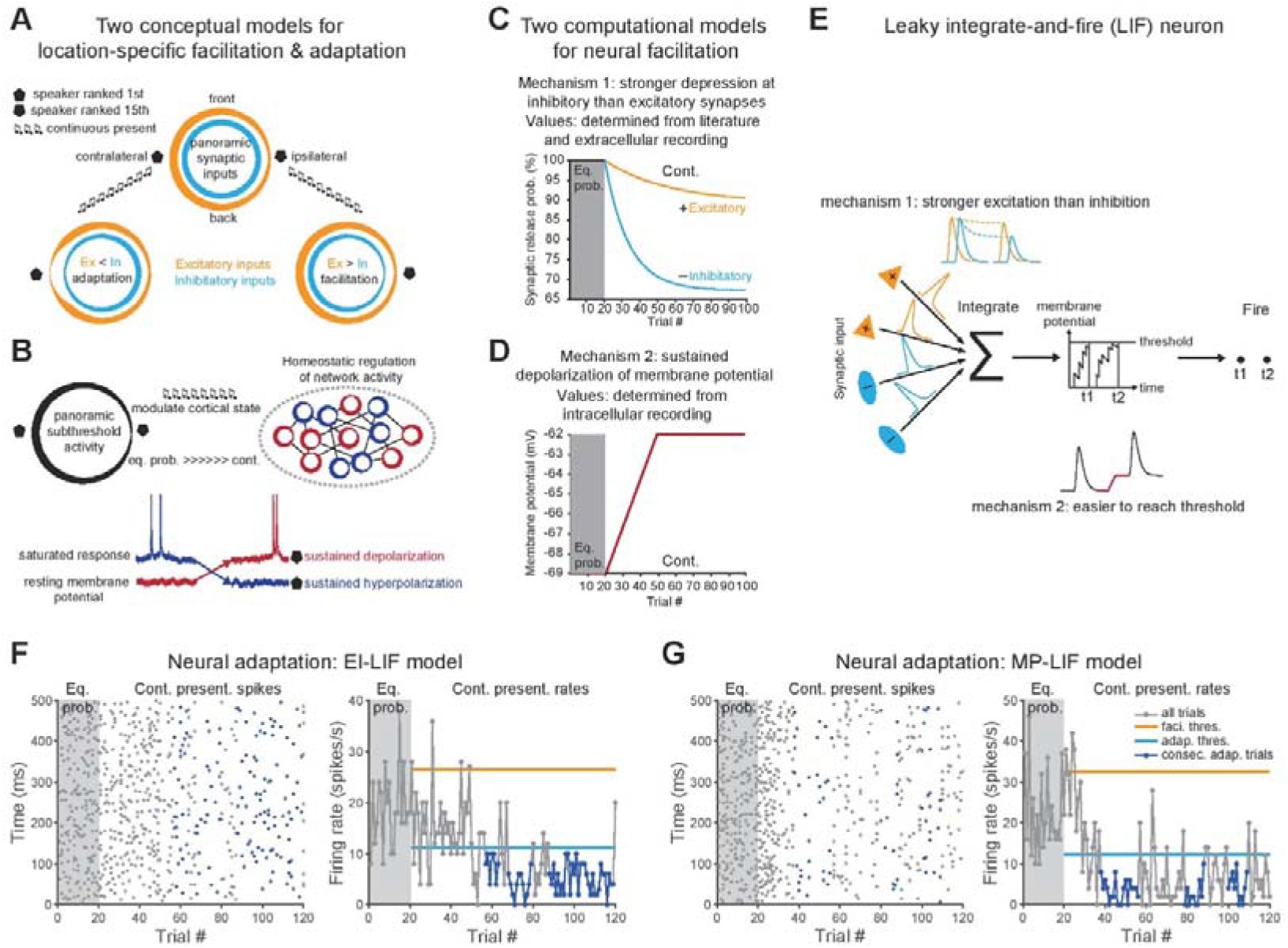
Explanations of EI-LIF and MP-LIF models. (A) A conceptual model for location-specific facilitation and adaptation. Orange and blue circles indicate excitatory and inhibitory synaptic inputs, respectively. The width of circles indicates the strength of synaptic inputs. Pentagon and upside-down pentagon shapes indicate 1st and 15th ranked speakers, respectively. Top, under equal-probability presentation mode, EI-LIF model neuron has a lower response at the ipsilateral location due to stronger inhibition (thick blue line) than excitation (thin orange line). Left, under continuous presentation mode, overall stronger inhibition than excitation would evoke neural adaptation at speaker ranked 1st (blue line is thicker than the orange line). Right, under continuous presentation mode, overall stronger excitation than inhibition would evoke neural facilitation at speaker ranked 15th (orange line is thicker than the blue line). (B) A conceptual model for location-specific sustained depolarization and hyperpolarization. Left, under equal-probability presentation mode, MP-LIF model neuron has panoramic subthreshold activity across all spatial locations but activity at the ipsilateral location is weakest (width of circle). Right, the neural activity of the network is homeostatically regulated: some neurons exhibit sustained depolarization (red circles) while others exhibit sustained hyperpolarization (blue circles). Black lines indicate the connections between two neurons. Bottom, under continuous presentation mode, sustained depolarization of weaker responses (red line) at the 15th-ranked speaker will be accompanied by sustained hyperpolarization of stronger responses (blue line) at the 1st-ranked speaker. (C) A computational model for location-specific facilitation. Gray shaded area (trial #1 to #20) indicates the equal-probability presentation mode. Under this presentation mode, no adaptation occurred for both excitatory and inhibitory synaptic inputs, thus the synaptic release probability is 100% for both inputs. Under the continuous presentation mode (trial #21 to #100), the inhibitory synapses (blue line) have a stronger adaptation amplitude (i.e., lower synaptic release probability) than the excitatory synapses (orange line). (D) A computational model for location-specific sustained depolarization. Under the equal-probability presentation mode, the membrane potential is stable at −69mV. Under the continuous presentation mode, membrane potential reached a stable level of −62mV which was 7mV depolarized after 30 trials. Those three parameters were obtained from our intra-cellular recordings. (E) Leaky integrate-and-fire (LIF) neuron model. LIF neuron integrates multiple excitatory (orange lines) and inhibitory (blue lines) synaptic inputs and fires a spike (t1, t2) whenever the membrane potential passes the threshold. Top, location-specific facilitation mechanism for the EI-LIF model neuron. Bottom, the other mechanism for the MP-LIF model neuron. (F) Neural adaptation simulated by EI-LIF model neuron. Left, spike raster plot of one session under equal-probability (shaded area) and continuous presentation mode. Blue dots indicate spikes belong to the long adaptation phase (i.e., at least five consecutive trials with firing rates lower than the adaptation threshold). Right, trial-by-trial firing rate from the same session. Blue dots and line indicate trails belong to the long adaptation phase. Thick orange and blue lines indicate the facilitation and adaptation thresholds, respectively. (G) Similar to (F) but for MP-LIF model neuron.

**Table S1.** Parameters and corresponding values used in EI-LIF and MP-LIF model neurons.

| Name | Symbol | Source | Value | Range |
| --- | --- | --- | --- | --- |
| Leaky integrate-and-fire (LIF, equation 1) |  |  |  |  |
| Time step | $\Delta t$ | This model | 0.0001s | fixed |
| Capacitance | $C$ | Wehr and Zador 2003 | $0.25 \cdot 10^{-9} \text{F}$ | fixed |
| Excitatory reversal potential | $E_e$ | Wehr and Zador 2003 | 0V | fixed |
| Inhibitory reversal potential | $E_i$ | Wehr and Zador 2003 | -0.085V | fixed |
| Leak conductance | $g_{rest}$ | Wehr and Zador 2003 | $25 \cdot 10^{-9} \text{S}$ | fixed |
| Resting potential | $E_{rest}$ | Experimental data | -0.069V | fixed (EI-LIF) |
| Spontaneous noise scale | $\sigma_s$ | This model | 0.01V | fixed |
| Gaussian noise | $\omega_n$ | This model | [-1: 1] | random |
| Threshold above resting potential | $V_{th}$ | Wehr and Zador 2003 | 0.02V | fixed |
| Excitatory and inhibitory conductance (equations 2 and 3) |  |  |  |  |
| Excitatory synaptic release probability | $P_{rel\_e}$ | This model | 1 | fixed (MP-LIF) |
| Inhibitory to excitatory ratio | $r$ | This model | 1 | fixed |
| Excitatory synaptic input number | $N_e$ | Wehr and Zador 2003 | 10 | fixed |
| Alpha function time constant | $\tau$ | Wehr and Zador 2003 | 0.005S | fixed |
| Conductance noise scale | $\sigma_c$ | This model | $2.5 \cdot 10^{-8} \text{S}$ | fixed |
| Inhibitory synaptic release probability | $P_{rel\_i}$ | This model | 1 | fixed (MP-LIF) |
| Inhibitory synaptic input number | $N_i$ | Wehr and Zador 2003 | 10 | fixed |
| Inhibitory to excitatory delay | $d$ | This model | 0S | fixed |
| Excitatory and inhibitory synaptic depression model (EI-LIF) (equations 4 and 5) |  |  |  |  |
| Excitatory synaptic depression amplitude | $A_e$ | This model | 0.003 | fixed |
| Excitatory synaptic time constant | $\tau_e$ | This model | 20S | fixed |
| Probability of target speaker | $P_s$ | Experimental data | 1/0.75/0.5<br>0.25/0.07 | fixed |
| Inhibitory synaptic depression amplitude | $A_i$ | This model | 0.025 | fixed |
| Inhibitory synaptic time constant | $\tau_i$ | This model | 10S | fixed |
| Stimulus count | $N_{SC}$ | Experimental data | 36 | fixed |
| Trial step | $T$ | This model | 1 | fixed |
| Number of trials in random mode | $T_r$ | This model | 20 | fixed |
| Number of trials in continuous mode | $T_c$ | This model | 300 | fixed |
| Membrane potential depolarization and hyperpolarization model (MP-LIF) (equations 6 and 7) |  |  |  |  |
| Dynamic resting membrane potential | $E_{rest\_MP}$ | Experimental data | -0.062V<br>-0.076V | max<br>min |
| Membrane potential polarization value | $M$ | Experimental data | $\pm 0.007 \text{V}$ | max, min |
| Number of trials to stabilize | $T_{th}$ | Experimental data | 30 | fixed |
| Spike threshold scale | $S$ | This model | 0.3 | fixed |
| Stimulus count | $N_{SC}$ | Experimental data | 30 | fixed |

